# CLARITY increases sensitivity and specificity of fluorescence immunostaining in long-term archived human brain tissue

**DOI:** 10.1101/2022.04.27.489700

**Authors:** Sarah Woelfle, Dhruva Deshpande, Simone Feldengut, Francesco Roselli, Karl Deisseroth, Jens Michaelis, Tobias M. Boeckers, Michael Schön

## Abstract

Immunohistochemistry on archival human brains is often limited because several conditions arise that complicate the use for high-resolution fluorescence microscopy. In this study, we developed a novel clearing approach for immunofluorescence-based analysis of perfusion- and immersion-fixed *post mortem* human brain tissue, termed hCLARITY. hCLARITY is optimized for specificity by reducing off-target labeling and yields very sensitive stainings in human brain sections allowing for super-resolution microscopy with unprecedented imaging of pre- and postsynaptic compartments. Moreover, hallmarks of the Alzheimer’s disease were preserved with hCLARITY, and importantly classical DAB or Nissl stainings are compatible with this protocol. hCLARITY is extremely versatile as demonstrated by the use of more than 30 well performing antibodies and allows for de- and subsequent re-staining of the same tissue section, which is important for multi-labelling approaches, e.g., in super-resolution microscopy. Taken together, hCLARITY enables research of the human brain with highest sensitivity and down to sub-diffraction resolution and therefore has enormous potential for the investigation of local morphological changes, e.g., in neurodegenerative diseases.

**Striking Image / Graphical abstract:** **Summary of the main advantages of hCLARITY.** CLARITY was applied on human *post mortem* brain tissue in direct comparison to untreated sections (minus CLARITY). Owing to removal of lipids during the clearing with SDS and the resulting reduction in light scattering, cleared sections were less opaque. The denaturing effect of the detergent SDS most likely results in better accessibility of certain epitopes. The major benefits of hCLARITY were found in an increased sensitivity and specificity of antibodies for neuronal cells. The adapted hCLARITY protocol is compatible with staining techniques for confocal microscopy, super-resolution microscopy, and light microscopy.

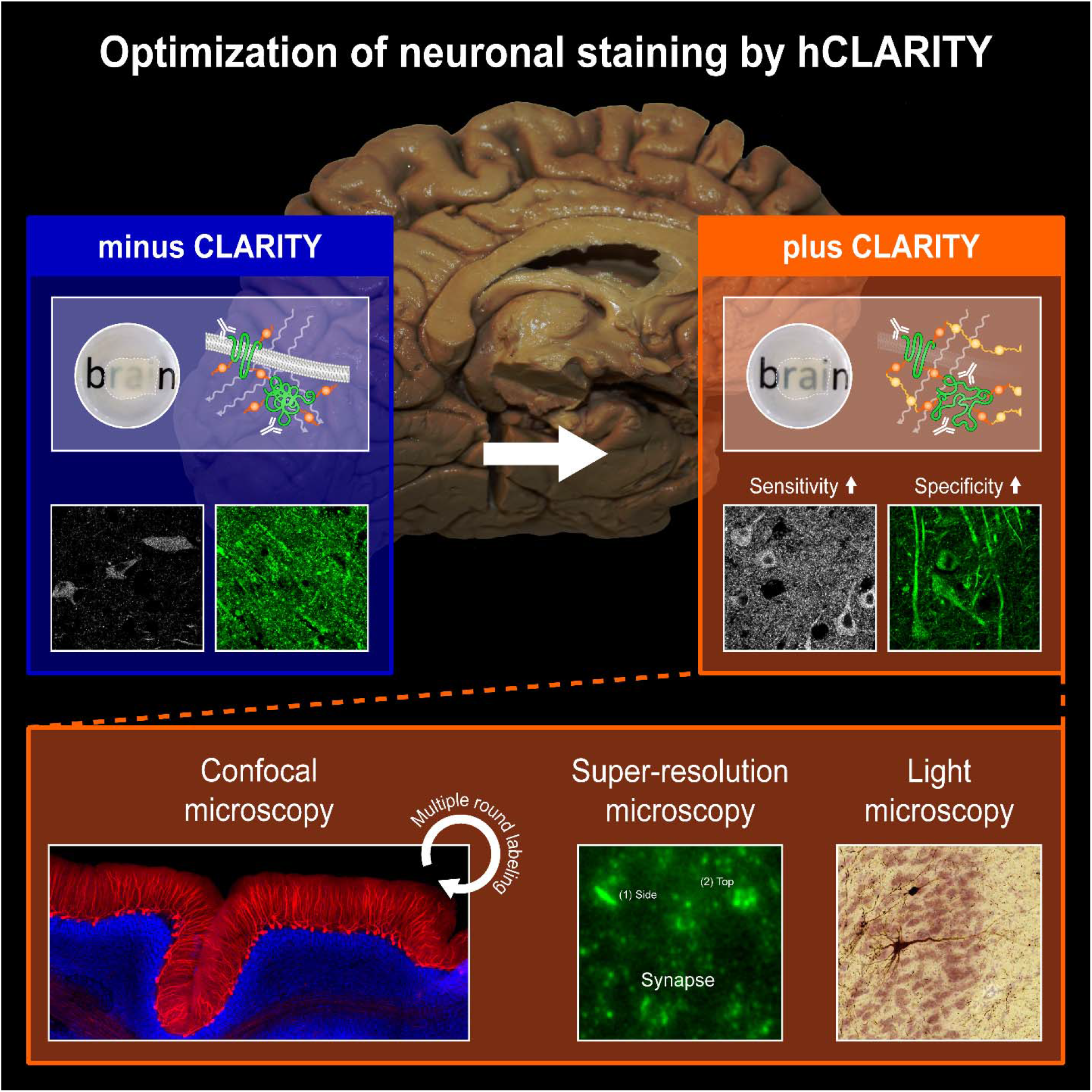

## 1. Introduction

*Post mortem* human brain tissue represents an indispensable source for the interrogation of molecules, structures, functions, and especially their changes during the course of neurodegenerative diseases. A major hindrance to quantifiable studies of the human brain is limited access to human brain tissue, although some research institutes have brain archives, and anatomists have access to a large number of donor tissue. In contrast to immunohistochemistry (IHC) on animal tissue, fixation conditions are usually not optimal, e.g., immersion rather than perfusion is mainly used as fixation method and, thus, the tissue is fixed only by diffusion. Finally, long-term archiving in formalin can reduce the immunogenicity of the contained proteins (Grillo et al., 2015). For these reasons, beyond classical non-immuno-based staining or chromogen-based staining of neurodegenerative protein deposits, it is difficult to image brain structures specifically and, most importantly, comparably. This, however, is the prerequisite for performing quantitative assessments.

One such application is the assessment of synapse loss in neurodegenerative diseases (Arendt, 2009; Clare, King, Wirenfeldt, & Vinters, 2010; de Wilde, Overk, Sijben, & Masliah, 2016), given the correlation between cognitive decline and synapse loss in Alzheimer’s disease (AD) (DeKosky & Scheff, 1990; Terry et al., 1991) and amyotrophic lateral sclerosis (ALS) (Henstridge et al., 2018). To date, synaptic densities have mostly been quantified only by microdensitometry (Leuba et al., 2008; X. Liu, Erikson, & Brun, 1996; Masliah et al., 2001; Masliah et al., 1994; Masliah, Terry, DeTeresa, & Hansen, 1989; Poirel et al., 2018; Terry et al., 1991), array tomography (Kay et al., 2013; Koffie et al., 2012), and electron microscopy (EM) (Dominguez-Alvaro, Montero-Crespo, Blazquez-Llorca, DeFelipe, & Alonso-Nanclares, 2019; Huttenlocher, 1979; Neuman et al., 2015; Scheff, DeKosky, & Price, 1990; Scheff, Price, Schmitt, & Mufson, 2006; Scheff, Price, Schmitt, Scheff, & Mufson, 2011). Disadvantages of these techniques are the low information content to a single synapse with densitometry, whereas 3D recordings with array tomography are laborious and only indirectly possible as serial recordings, and immunogold labeling with EM is highly demanding. Most importantly, each of these methods usually requires freshly fixed tissue. Thus, model systems are favored as surrogates under more controllable conditions, e.g., mouse models (Di et al., 2016; Neuman et al., 2015) or induced pluripotent stem cells and their neuronal derivatives (Penney, Ralvenius, & Tsai, 2020). However, such models do not faithfully represent all conditions in the human organism as the ‘original’, and, in neurodegenerative diseases of the aging human brain, model systems do not reach the advanced age of brains from deceased individuals. Finally, clinical symptomatology and histopathology can only be directly matched with human brain tissue.

The clearing technique CLARITY was invented with the major goal of enabling coherent 3D labeling and microscopy of intact tissues or whole organs, thereby making laborious and error-prone serial sectioning approaches superfluous (Chung et al., 2013). In addition, immunofluorescence staining can be performed after CLARITY, which overcomes the limitations inherent in non-immuno-based and chromogen-based techniques. In the here applied and adapted CLARITY protocol, clearing was achieved by immersing the hydrogel-stabilized tissue in a 4% sodium dodecyl sulfate (SDS) solution.

Since SDS is known as an antigen retrieval agent (Brown et al., 1996) and speculation exists about its benefits on immunostaining after CLARITY (A. K. Liu et al., 2016), we referred to the study of Brown et *al.* (Brown et al., 1996) and, thus, we further assessed the detergent-related effects on the accessibility of epitopes in *post mortem* human brain tissue. We used perfusion-fixed brains from the gross anatomy course and immersion-fixed brains from the tissue bank collection at Ulm University. First, by performing confocal microscopy, we evaluated the staining quality of a selection of neuronal and synaptic markers in direct comparison to non-cleared sections. Here, besides the reported increase in tissue translucence owing to lipid removal via the CLARITY method, we found that some antibodies stained more sensitively and specifically, i.e., that staining of target structures resulted in a stronger signal (greater sensitivity) and displayed less background (greater specificity). Second, whereas prior studies have demonstrated the applicability of CLARITY to human neural tissue using only a few antibodies or without a systematic investigation of their benefits (Morawski et al., 2018; Phillips et al., 2016), we tested a large panel of neuronal, non-neuronal, and Alzheimer’s pathology markers. We established an ‘antibody toolbox’ with 30 markers as a ready-to-use panel saving time and costs for antibody testing. In addition, by stabilizing the tissue with acrylamide, CLARITY offers easy detergent-based de-staining and subsequent re-staining rounds (multiple round labeling). Finally, we show that passive CLARITY is also compatible with super-resolution microscopy (stimulated emission depletion microscopy (STED), direct stochastical optical reconstruction microscopy (dSTORM)) for imaging human cortical synapses in perfusion-fixed tissue. These techniques were mainly applied to perfusion-fixed tissue from the gross anatomy course at Ulm University, a source probably available in many anatomical institutions at universities. This fixation technique proved to be superior for immunofluorescence staining, which is in line with a recently published study (McFadden et al., 2019). Exploiting this source in combination with the here-presented tools will provide a broader basis for future studies on human tissue. We termed our adapted passive CLARITY protocol hCLARITY (‘human CLARITY’).

## 2. Materials and Methods

### 2.1 Human brain samples

Perfusion-fixed human *post mortem* brains (cases with a p (perfusion) code, main cases were p1 and p2 for most of the experiments; cases p1-p2 in Table 1) were obtained from permanent body donors of the gross anatomy course organized by the Institute for Anatomy and Cell Biology, Ulm University. All included donors provided their informed and written consent for permanent body donation. This study was approved by the ethics committee of Ulm University. After death, the cadavers were fixed with a solution according to Tutsch by perfusion via the femoral artery (per body donor: 8 l ethanol, 3 l glycerol 86% (Carl Roth, Karlsruhe, Germany), 0.6 l lysoformin (Lysoform Dr. Hans Rosemann GmbH, Berlin, Germany), 0.6 l formaldehyde 30% (Carl Roth, Karlsruhe, Germany), filled up to 15 l with water). More precisely, the femoral artery was opened longitudinally and 1.5 l of the fixans was introduced distally. Then, the distal part of the artery was closed, and 11.5 l of the fixans were infused proximally. The artery was sutured closed. Thereafter, the cadavers were stored for a minimum of four months for post-fixation and were continuously moistened with 5% formaldehyde solution. At the end of each anatomy course, brains were collected and immersed in a 1% aqueous solution of formaldehyde. Until usage, brains were stored in a dark room.

**Table 1.** Demographics and clinical-neuropathological diagnoses of the n = 6 cases studied.

| Case | Age (yrs) | Sex | NFT stage | Aβ phase | PD stage | PMI (h) | Fixation year | Origin | | Cause of death |
| --- | --- | --- | --- | --- | --- | --- | --- | --- | --- | --- |
|  |  |  |  |  |  |  |  | Anatomy course | Tissue bank |  |
| p1 | 71 | M | I | 0 | 0 | 72 | 2019 | x |  | Cardiogenic shock |
| p2 | 74 | M | I | 3 | 0 | 24 | 2017 | x |  | Decompensated heart insufficiency |
| i1 | 49 | F | 0 | 0 | 0 | N.a. | 2016 |  | x | N.a. |
| i2 | 82 | F | II | 1 | 0 | 71 | 2015 |  | x | Stroke |
| i3 | 80 | M | III | 0 | 0 | 39 | 2003 |  | x | Malignant neoplasm |
| i4 | 78 | M | I | 0 | 0 | 56 | 2003 |  | x | COPD |

**Abbreviations:** yrs = years; NFT = neurofibrillary tangle; Aβ = amyloid-β ; PD = Parkinson's disease; PMI = *post mortem* interval; h = hour; P = perfusion fixation; I = immersion fixation; F = female; M = male; n.a. = not available; COPD = chronic obstructive pulmonary disease

Centimeter-thick tissue blocks of the desired brain regions were excised and cut on a vibratome (Microm HM 650 V, Thermo Scientific/Microm, Walldorf, Germany and VT1200S vibratome, Leica, Wetzlar, Germany) into 100 µm thick sections. All cases used for evaluating the effects of CLARITY on immunostaining are described in Table 1. Cases p1 and p2 were also used to assess the performance of the majority of antibodies (Table 2). If no code for the case is indicated, representative images were derived from one of ten additional perfusion-fixed cases. For comparison with the perfusion-fixed anatomy course cases, immersion-fixed brains from the tissue bank collection at Ulm University were used in selective experiments (cases i1-i4 in Table 1), which were fixed in a 4% aqueous solution of formaldehyde (H. Braak & Del Tredici, 2015).

**Table 2.**
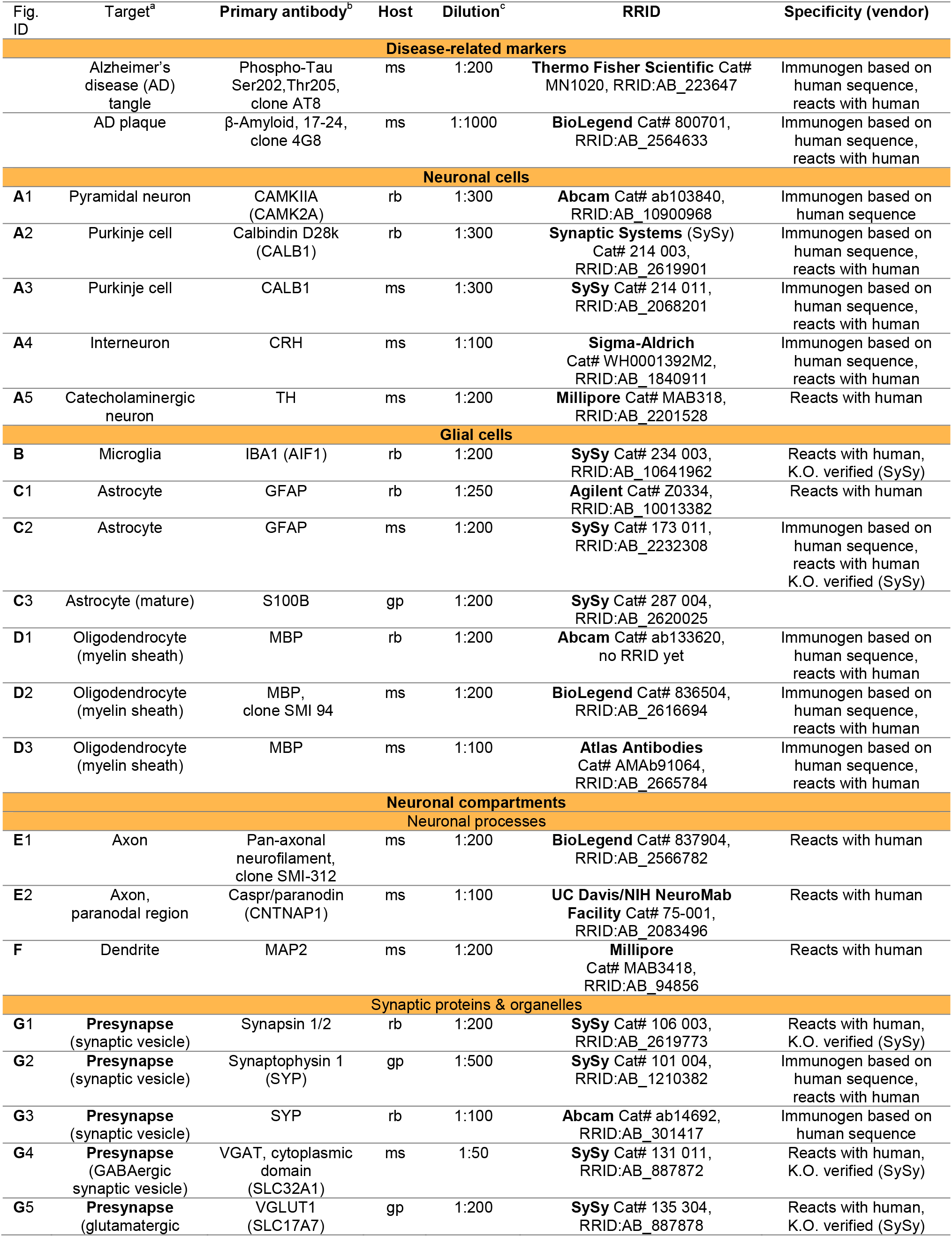
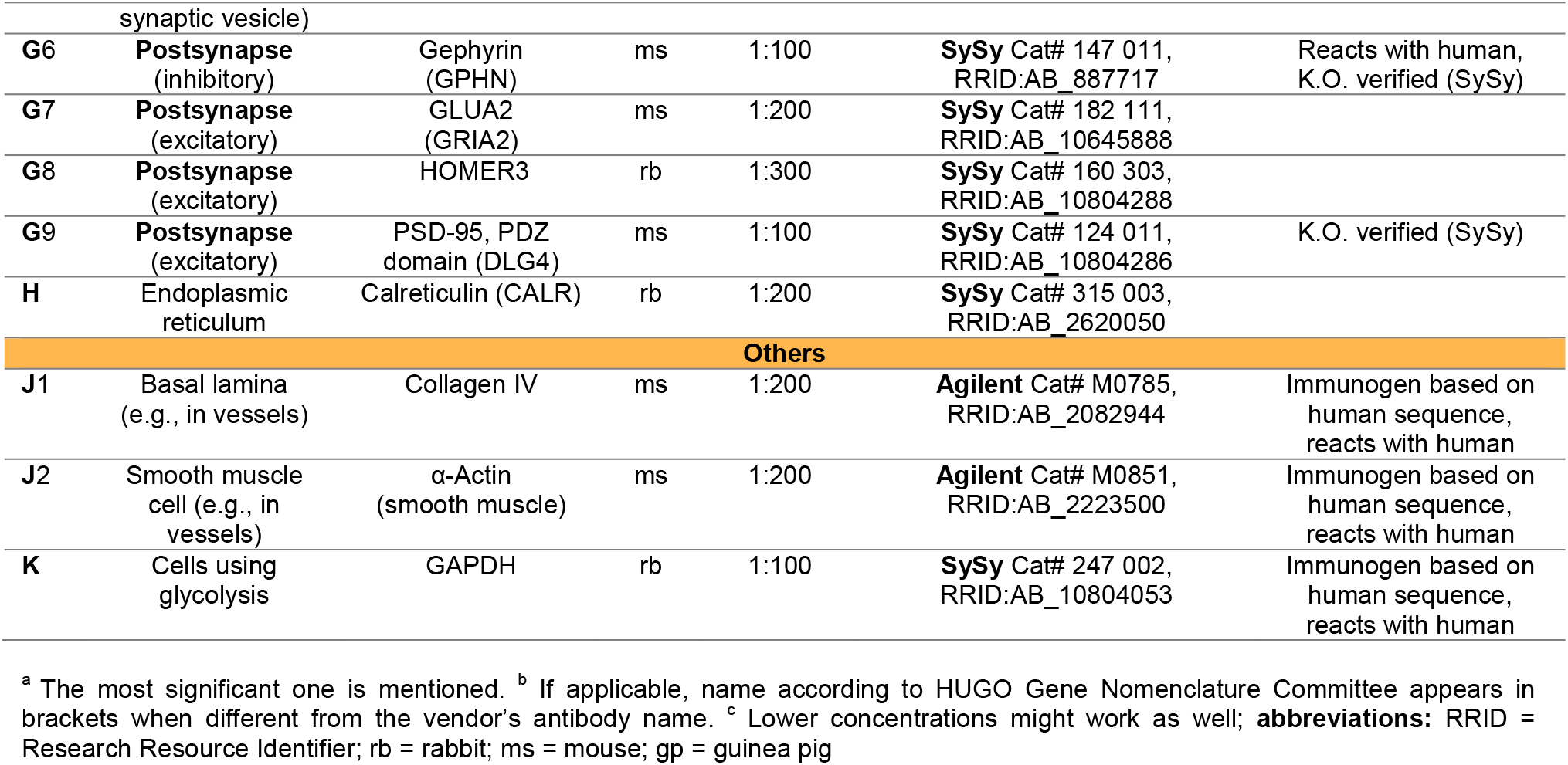
Detailed information on the 30 primary antibodies used.

### 2.2 CLARITY

CLARITY procedure and solutions were based on previously published protocols (Chung et al., 2013; Tomer, Ye, Hsueh, & Deisseroth, 2014). Hydrogel was prepared separately as solution I and II: Solution I was prepared by dissolving 8 g (16% (wt/vol)) paraformaldehyde (Sigma-Aldrich, Steinheim, Germany) (PFA, 4% (vol/vol) final concentration) in 50 ml Dulbecco’s phosphate-buffered saline (DPBS) (Thermo Fisher Scientific, Waltham, MA). Solvation was achieved by adding several drops NaOH and by stirring at 60 °C. For solution II, 20 ml of 40% acrylamide (Bio-Rad, Munich, Germany) (4% (vol/vol) final concentration), 20 ml 10x DPBS, and 110 ml dH_2_O were mixed. After combining solution I and II, the pH of the hydrogel was adjusted to 7.35-7.45 and the 200 ml were filtered. Immediately before immersing the sections in hydrogel, 0.25% (wt/vol) of the thermal initiator VA-044 (Wako Chemicals, Neuss, Germany) was admixed. Incubation in hydrogel took place for 5 days at 4 °C.

To enable polymerization of the hydrogel, oxygen was removed by vacuumizing the samples for 15 min at 0-20 mbar and subsequently adding nitrogen in a desiccation chamber. Then, polymerization was started by placing the samples in an incubator for 3 hours at 37 °C. The hydrogel was replaced after the first 1.5 hours by PBS to prevent the solidification of hydrogel on the sections’ surfaces. After polymerization, passive clearing was initiated by changing the sections to clearing solution (4% (wt/vol) sodium dodecyl sulfate (SDS) (Serva, Heidelberg, Germany), 200 mM boric acid (Merck, Darmstadt, Germany); pH 7.0-7.4). To remove any residual hydrogel, the clearing solution was exchanged on the first and second day after starting the passive clearing. Clearing of the sections took place for 7 days at 37 °C under gentle agitation on a shaking table.

### 2.3 Immunohistochemistry

#### Fluorescence-based immunolabeling for confocal microscopy

At the conclusion of the passive clearing process, tissue sections were rinsed three times within 24 h with PBS plus Triton X-100 (PBST; 0.1% (vol/vol) Triton X-100 (Roche, Mannheim, Germany)) under gentle agitation at room temperature. Prior to immunostaining, heat-induced epitope retrieval was performed with 0.1 M citrate buffer (pH 6.0). Briefly, sections were transferred into glass cuvettes filled with citrate buffer and boiled for 10 min. Afterwards, sections were incubated in blocking solution (3% (wt/vol) bovine serum albumin (BSA) (Bio Froxx, Einhausen, Germany) plus 0.3% (vol/vol) Triton X-100 in PBS) for 3 h under gentle agitation. Primary antibodies were diluted in blocking solution according to the protocol in Table 2 and were incubated with the sections for 48 hours at 4 °C under gentle agitation. Subsequently, rinsing with PBS (1x 1 h, 2x 30 min) was followed by staining with Alexa Fluor-conjugated secondary antibodies (Invitrogen, Waltham, MA and Jackson ImmunoResearch Laboratories, Inc., Ely, UK, 1:200 in blocking solution) for 3 h at room temperature. Thereafter, the first 1 h rinsing step included 4’,6-diamidino-2-phenylindole (DAPI (Carl Roth, Karlsruhe, Germany), stock solution 10 mg/ml, diluted 1:10,000 in PBS), followed by 2x 30 min washing with PBS. The aforementioned incubation steps were performed under gentle agitation at room temperature.

Finally, tissue sections were mounted in ProLong Gold antifade mountant (Invitrogen) or self-made refractive index matching solution (RIMS; according to (Yang et al., 2014)) and were cover-slipped. Sections for re-staining were mounted in aqueous mountants only (RIMS or SlowFade Gold antifade mountant, Invitrogen). For experiments with a comparison of plus and minus CLARITY staining results, ProLong Gold antifade mountant was consistently applied. In each experiment, additional tissue sections were incubated with the secondary antibodies only (negative control: omission of primary antibody) to control for unspecific binding. At least one representative staining of each antibody in Table 2 is shown in the main figures or in the figure supplements at the end of this manuscript.

#### Re-staining after confocal microscopy

Tissue sections mounted in aqueous mounting media (RIMS or SlowFade Gold antifade mountant) were subjected to a second staining round. By immersion of slides in PBS, coverslips were removed and sections could be carefully detached from the slide. For de-staining, sections were incubated in clearing solution at 60 °C for 24 h. Subsequently, sections were shortly rinsed in PBS, followed by a longer washing step in PBST. For re-staining, see section “Fluorescence-based immunolabeling for confocal microscopy”, starting from the heat-induced epitope retrieval step. Control sections were mounted without re-staining and imaged with a confocal microscope.

#### Fluorescence-based labeling for stimulated emission depletion (STED) microscopy

Cleared 50 µm thick sections were immunostained as described above. GLUA2 and MAP2 antibodies were diluted as indicated in Table 2 (G7 & F), the CAMKIIA antibody (A1) was diluted 1:200. Synaptic staining was performed with a synaptophysin (1:300, Synaptic Systems, Göttingen, Germany, G2 in Table 2) and a GLUA1 antibody (1:200, Synaptic Systems, Cat. No. 182011, RRID: AB_2113443). Instead of Alexa Fluor-coupled secondary antibodies, ATTO dyes (Sigma-Aldrich; ATTO647N, Cat. No. 50185 and ATTO594, Cat. No. 77671 and Cat. No. 76085) were used for labeling. When commercial dye-labeled antibodies were not available, antibodies were covalently labeled in-house. Donkey anti-guinea pig secondary antibodies (Jackson ImmunoResearch Laboratories, Inc.; RRID: AB_2340442) were covalently labeled with ATTO647N succinimidyl ester or ATTO594 succinimidyl ester dye (ATTO-TEC, Siegen, Germany) using manufacturer’s protocol. Briefly, two molar excess of reactive dye was incubated with unconjugated antibodies in a 0.1 M sodium bicarbonate (Honeywell Fluka, Seelze, Germany) solution for 30 min under agitation. The unbound fluorophores were removed by gel filtration on a NAP-5 column (Sephadex-25 medium, GE Healthcare, Munich, Germany). The final antibody concentration and degree of labelling was analyzed on a spectrophotometer (NanoDrop 2000 UV-Vis; Thermo Scientific). Slides and coverslips were cleaned thoroughly and sections were finally mounted in 97% (vol/vol) 2,2’-thiodiethanol (TDE, Sigma-Aldrich) in PBS (pH 7.5) for refractive index matching.

#### Fluorescence-based labeling for direct stochastic optical reconstruction microscopy (dSTORM)

Cleared 50 µm thick sections were immunostained as described above. Synaptic staining was performed with the same synaptophysin and GLUA1 primary antibodies (dilution each 1:200), as described for STED microscopy. Alexa Fluor-conjugated secondary antibodies (1:350, Invitrogen, Alexa Fluor 647, Cat. No. A21450 and Alexa Fluor 532, Cat. No. 11002) were applied 90 min for staining, followed by three rinses à 30 min with PBS. To facilitate photo switching of dyes, samples were mounted in a special imaging buffer (100 mM cysteamine, 400 U/ml of catalase, 100 U/ml glucose oxidase, and 40 mg/ml of glucose; all Sigma-Aldrich).

#### Chromogen-based labeling (3,3’-diaminobenzidine, DAB)

Neuropathological stages for Alzheimer’s disease (AD) of cases from the gross anatomy course are provided in Table 1. For neurofibrillary tangle (NFT) staging, the required tissue blocks according to (H. Braak, Alafuzoff, Arzberger, Kretzschmar, & Del Tredici, 2006) were excised. For determination of amyloid-β (Aβ) phases, tissue blocks were assessed as described previously (Thal et al., 2000). Despite the selection of low-stage cases, the cerebellum was additionally assessed to diagnose Aβ phase 5 (Thal, Rüb, Orantes, & Braak, 2002). Parkinson’s disease (PD) neuropathological staging was performed according to (H. Braak et al., 2003). The tissue blocks were cut into 100 µm thick sections with a vibratome and on the same day, or the day after, immunolabeling was started as previously described in detail (H. Braak, Thal, Ghebremedhin, & Del Tredici, 2011). Briefly, after blocking endogenous peroxidase, sections for detection of Aβ deposits and Lewy pathology (α-synuclein) underwent antigen retrieval for 3 min with 100% formic acid (Appli Chem, Darmstadt, Germany), whereas detection of hyperphosphorylated tau was performed without retrieval. After blocking with BSA, sections were incubated overnight with an antibody directed against phospho-tau (clone AT8, 1:2000, Thermo Fisher Scientific, Cat. No. MN1020), β-amyloid (clone 4G8, 1:5000, BioLegend, Cat. No. 800701) or α-synuclein (1:2000, BD Transduction Laboratories, Cat. No. 610787, RRID: AB_398108). Following incubation with a biotinylated secondary antibody (1:200, Vector Laboratories, Burlingame, CA) and signal amplification via the ABC kit (Vectastain Elite ABC-HRP Kit, PK6100; Vector Laboratories), reactions were finally visualized with chromogen 3,3’-diaminobenzidine (DAB, CN75, Carl Roth). Positive (section with abundant pathology) and negative (omission of primary antibody) controls were included. Sections were dehydrated in an ascending alcohol series, cleared in Histo-Clear (National Diagnostics, Atlanta, GA) or xylene, and mounted in Histomount (National Diagnostics) or Entellan New (Merck) (see also (Feldengut, Del Tredici, & Braak, 2013)). Neuropathological stages for cases from the tissue bank are also shown in Table 1.

#### DAB-based labeling and Nissl staining after CLARITY

The staining started after the one-day washing step (three exchanges of PBST, see above). The same protocol as described above was used for subsequent detection of phospho-tau, counterstaining with Darrow red (Sigma-Aldrich, Cat. No. 211885) was performed for detection of basophilic Nissl substance (Heiko Braak, 1980; H. Braak, Braak, Ohm, & Bohl, 1988). For CAMKIIA staining (1:300, A1 in Table 2), no additional retrieval was performed, the goat anti-rabbit biotinylated secondary antibody was also purchased from Vector Laboratories (Cat. No. BA-1000).

### 2.4 Image acquisition

#### Confocal microscopy

Images were captured with a Leica SPE confocal microscope (Leica, Germany) using laser excitation at 635 nm, 561 nm, 488 nm, and 405 nm in that order. Samples were viewed with a 40x oil objective (ACS APO; numerical aperture (NA) 1.15; free working distance (WD) 270 µm) or a 63x oil objective (ACS APO; NA 1.30; WD 160 µm). Two overview scans were acquired using a 10x dry objective (ACS APO; NA 0.3; WD 3 mm) and a 20x multi-immersion objective (ACS APO; NA 0.60; WD 200 µm) with oil. For z-stacks, settings were kept constant in individual z-planes.

#### STED microscopy

Images were acquired on a custom-built dual-color STED microscope with a 100x oil objective (HCX PL APO 100x/1.40-0.70 oil CS, Leica). A detailed description of the setup can be found in (Osseforth, Moffitt, Schermelleh, & Michaelis, 2014). The system enables switching between confocal (excitation at 568 nm and 633 nm) and STED (additional depletion beams at 715 nm and 750 nm) mode. The excitation lasers were used at a power of 0.6-1 µW and the depletion laser power was ∼1.8 mW. Initially, confocal images were acquired at a pixel size of 50 nm or 100 nm, followed by the corresponding STED images at a pixel size of 20 nm. The pixel dwell time was 200 µs (confocal mode) and 300 µs (STED mode).

#### dSTORM imaging

Imaging was performed on a custom-built widefield setup with a 60x oil immersion objective (60x APO TIRF, NA 1.49 Oil, Nikon, Tokyo, Japan). Samples were imaged using 405 nm activation laser and excitation lasers 647 nm or 532 nm; fluorescence was detected with an EMCCD camera (Andor Technology, Belfast, UK). The setup construction is detailed in (Deshpande et al., 2019; Schoen et al., 2015). First, widefield images were captured for both channels. Then, for each channel for image reconstruction, ca. 30,000 frames were recorded consecutively at a 20 ms exposure time. Single molecule localizations were determined using custom algorithms written in MATLAB (MathWorks, Natick, MA) as described before (Schoen et al., 2015) (see 2.5).

#### Conventional light microscopy

DAB-based CAMKIIA staining and the Nissl staining were imaged with the color camera of a Biorevo BZ-X810 microscope (Keyence, Neu-Isenburg, Germany) using a 100x oil objective (CFl Plan Apo λ 100XH; NA 1.45; WD 0.13 mm) and a 20x objective (CFl Plan Apo λ 20x; NA 0.75; WD 1 mm). Full focus of z-stacks was obtained with the BZ-X800 Analyzer (Keyence) software and exported as TIFF images. Additional DAB sections were acquired with a BX61 microscope (Olympus Optical, Tokyo, Japan). The single images from all layers of the stack were merged after the so-called EFI (extended focal imaging) algorithm extracted those features of the images with the sharpest contrast (Cell D Imaging Software (Olympus)).

### 2.5 Image Processing

#### Confocal microscopy

For comparison of plus and minus CLARITY sections, individual z-stack planes were extracted using the Leica Application Suite X (LAS-X) software, and brightness/contrast were adjusted identically with Imaris (Oxford Instruments, Abingdon, UK) software. Maximum intensity projections, 3D renderings, and video material were also created in Imaris software. The described brightness/contrast adjustments were performed on entire z-stacks; no individual adjustments of single z-planes were done. For most cases, a Gaussian filter with default filter width in Imaris was applied prior to brightness/contrast adjustment.

#### STED microscopy

Confocal and STED images were brightness/contrast-adjusted in Fiji (ImageJ, NIH, Bethesda, MD) and a Gaussian blur with σ = 1.0 was applied for visualization. Intensity profiles were created on raw data using the ‘plot profile’ command. Z-stack stabilization of confocal and STED stacks (Figure 7-figure supplement 1) was done with the Huygens Object Stabilizer (SVI, Hilversum, The Netherlands) using identical processing steps.

#### dSTORM reconstruction

Centre-of-mass fitting was done on the FIRESTORM software (previously described in (Schoen et al., 2015) and (Deshpande et al., 2019)). The localizations were filtered based on the intensity, full width at half maximum (FWHM), and symmetry distribution. A redundant cross-correlation algorithm was implemented to correct the xy-drift (Wang et al., 2014). The chromatic aberration between channels was corrected by using a pre-determined transformation function generated from imaging Tetraspeck beads. Final images were rendered with a 10 nm pixel size, weighted by intensity of individual localizations and the two channels were combined to have two-color super-resolved images. Analysis was always conducted on raw data; the processed image (Gauss filter) was used only for visualization purposes.

#### Conventional light microscopy

TIFF files were opened in Fiji (ImageJ) and contrast/brightness were adjusted. Adjustments were performed equally for plus and minus CLARITY sections.

### 2.6 Data analysis

Stack intensity profiles and line profiles were exported from LAS-X and Fiji (ImageJ), respectively, as Excel (Microsoft Corporation, Redmond, WA) files. Final graphs were created in GraphPad Prism (GraphPad Software, La Jolla, CA). FWHM was calculated on raw data using the non-linear fit command (Gauss function) of OriginPro (OriginLab, Northampton, MA). Final figures were set up in Adobe Illustrator (Adobe, San José, CA).

### 2.7 Western blot with human Protein Medley

50 µg of human cerebral cortex lysate (Protein Medley, Takara, Kusatsu, Japan, Cat. No. 635323) were diluted and prepared according to manufacturer’s instructions (protocol PT1602-1). The sample and a protein ladder (Spectra Multicolor Broad Range, Thermo Scientific) were loaded on a 4-15 % precast protein gel (Bio-Rad, Cat. No. 4561085). Western blotting was performed according to standard protocols, the GLUA2 antibody (Synaptic Systems, G7 in Table 2) was diluted 1:500 and β-actin was used as loading control (1:10,000, Sigma-Aldrich, Cat. No. A5316, RRID: AB_476743). An horseradish peroxidase (HRP)-coupled rabbit anti-mouse antibody (Dako, Glostrup, Denmark) in combination with Clarity Western ECL Substrate (Bio-Rad) were used for signal detection according to manufacturer’s instructions.

## 3. Results

As the results of this study include a variety of methodologies, a summary of the here applied and adapted CLARITY method for long-term formalin-fixed human brain tissue, referred to as hCLARITY, and its possible applications are depicted in Figure 1. With hCLARITY, human brain tissue fixed with different methods (immersion-fixed and perfusion-fixed, indicated with i or p in front of the case number (see Table 1 for further details)) or for different time periods can be utilized. Sections are either cut directly from the fixed tissue or from embedded tissue blocks. The sections are then stabilized by incubating them in a monomeric acrylamide-containing solution followed by polymerization introducing a hydrogel of polymeric acrylamide that binds to the protein-paraformaldehyde network. Subsequent detergent treatment on a shaking table at 37 °C, i.e., passive clearing, removes the biolipids that cause light scattering and a penetration barrier for the diffusion of macromolecules. hCLARITY is compatible with classical histological staining methods, such as Nissl, or chromogen-based IHC with 3,3’-diaminobenzidine (DAB) that is routinely used in neuropathology. In particular, hCLARITY is suitable for a broad variety of fluorescence-based imaging techniques, even with repetitive rounds of detergent based de-staining and subsequent re-staining (multiple round labeling). Below, we explain the results and benefits of our applications in greater detail.

**Figure 1.**
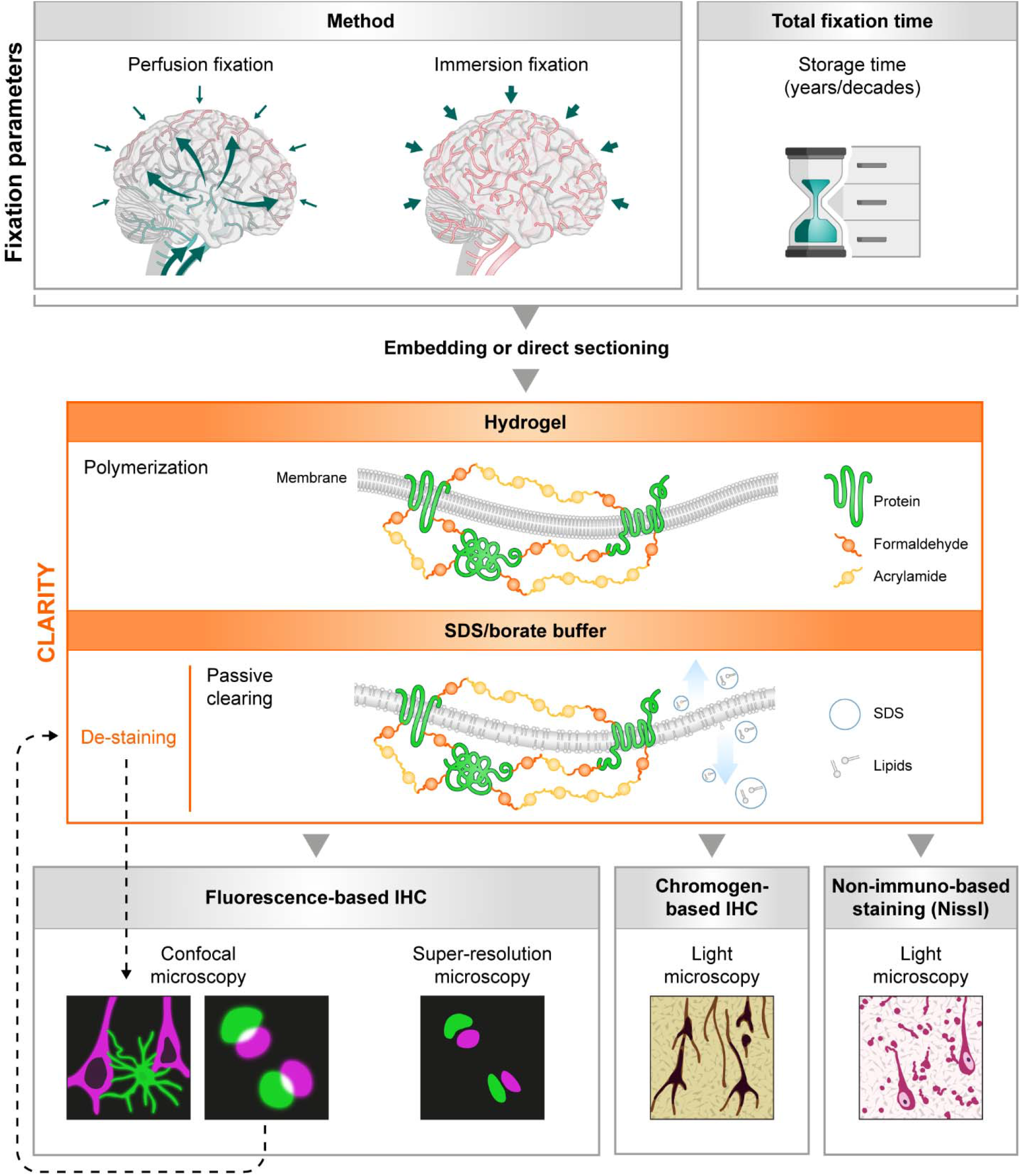
Flowchart of compatible staining and imaging techniques for CLARITY-treated *post mortem* human brain tissue. Independent of type of fixation, e.g., perfusion-fixed *versus* exclusively immersion-fixed, or total fixation time, tissue samples can be embedded or cut directly on a vibratome. After CLARITY treatment, consisting of hydrogel embedding and polymerization with a consecutive SDS-based clearing step, fluorescence-or chromogen (e.g., 3,3’-diaminobenzidine (DAB))-based staining with a variety of cellular and synaptic markers can be performed. In addition, non-immuno-based Nissl staining is also compatible with CLARITY. Imaging can be performed with fluorescence microscopy (confocal or super-resolution microscopy) or with conventional light microscopes in case of DAB and Nissl staining. Primary and secondary antibodies as well as DAPI can be de-stained with the clearing solution, allowing for multiple staining rounds.

### 3.1 Increased sensitivity of immunostaining after CLARITY

In two independent experimental rounds (n = two technical replicates), sections from the superior frontal gyrus (Brodmann area 9 (BA9)) underwent either CLARITY after cutting (= plus CLARITY) or they were cut shortly before immunolabeling (= minus CLARITY). For the IHC protocol, all sections were processed equally and in parallel.

First, we aimed to compare the staining quality of antibodies directed against neuronal cells starting with a calcium/calmodulin-dependent protein kinase II alpha (CAMKIIA) antibody. CAMKIIA is not exclusively expressed in neuronal cells but is also part of the postsynaptic density (PSD) (X. B. Liu & Murray, 2012). In the two perfusion-fixed cases, we found an overall higher mean intensity for plus CLARITY sections when viewed with identical parameters at a confocal microscope than minus CLARITY sections. This finding could be followed throughout a z-stack with z = 15 µm (Figure 2a). Of note, while the single planes extracted from the z-stack still showed positive labeling of neurons and their processes in minus CLARITY sections, the intensity of punctate structures, presumably highlighting PSDs, was clearly enhanced in plus CLARITY sections. For plane I of the z-stack, e.g., the mean intensity was increased approximately by a factor of 2 in cleared compared to non-cleared sections. Similar tendencies could be observed for the immersion-fixed cases with a total fixation time of 18 years (Figure 2-figure supplement 1a: mean intensity position I: factor 1.6 higher) and 5 years (Figure 2-figure supplement 1b: mean intensity position I: factor 2 higher). Importantly, the negative controls of cleared sections showed comparable background levels to non-cleared ones. For the remaining cases (i2, i3, and p2), CAMKIIA mean intensity in cleared compared to non-cleared tissue sections was on average twice as high (cases i2 and i3) or nearly by a factor three higher (case p2).

**Figure 2.**
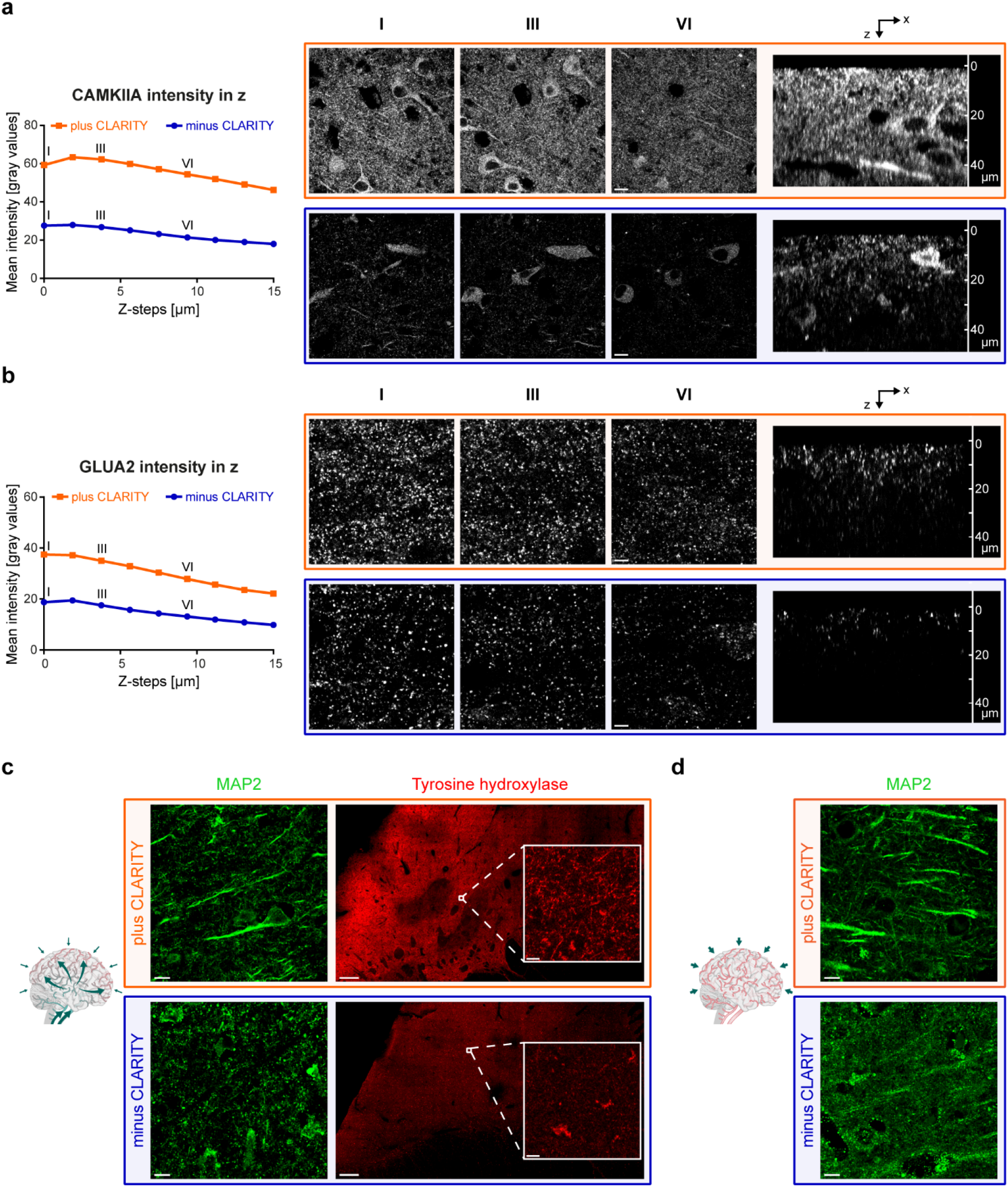
Increased sensitivity and specificity of antibodies applied on CLARITY-processed sections. **(a)** Cleared (orange) and non-cleared (blue) frontal cortex sections from case p1 were stained for CAMKIIA and imaged 15 µm deep in 1.8 µm steps using a confocal microscope. Mean intensities of each z-step were exported with LAS-X software from two stacks (cortex layer V), and the mean was plotted in dependency of the z-level. Z-level I (z = 0 µm), III (z = 3.6 µm), and VI (z = 9 µm) are provided as single images. A side view (xz scan) is shown on the far right. Scale bars, 10 µm. **(b)** Mean intensity of GLUA2 staining on case p1 was plotted in dependency of the z-level. Acquisition strategy is the same as outlined in (a). Scale bars, 5 µm. **(c)** MAP2 (frontal cortex, case p2) and tyrosine hydroxylase (caudate nucleus, case p2) labeling in perfusion-fixed cases plus (orange) and minus (blue) CLARITY treatment. Scale bars, 500 µm (TH), 10 µm (MAP2, TH magnification). **(d)** MAP2 labeling plus (orange) and minus (blue) CLARITY treatment in immersion-fixed brains with a comparable total fixation time as (c) (frontal cortex, case i1). Scale bars, 10 µm. See also Figure 2-figure supplement 1 and corresponding source data files for both figures.

To compare plus and minus CLARITY results in a laser power-independent setting, we performed traditional DAB-based CAMKIIA staining on cleared and non-cleared sections. Not only were CLARITY-treated sections compatible with subsequent DAB staining, but again more intense labeling for cleared sections in comparison to minus CLARITY samples was observed (Figure 2-figure supplement 1c).

Given the difficulty associated with staining postsynaptic structures, we then tried to see whether CLARITY improved staining quality of pre- and postsynaptic markers. In the perfusion-fixed cases, we found an overall higher mean intensity of glutamate receptor subunit 2 (GLUA2) throughout a stack with z = 15 µm, when viewed with identical parameters at a confocal microscope as compared to minus CLARITY sections. This became clearly visible in the xz scan (Figure 2b). For immersion-fixed cases, no generally applicable tendency was observed; in some cases, cleared sections revealed a finer and more homogenous punctate labeling (data not shown). Especially in the context of AD-related synaptic decline, synaptophysin (SYP) has been a popular marker showing robust results in immunofluorescence on human *post mortem* tissue, see, e.g., (Henstridge et al., 2018; Masliah et al., 1989). Here, we did not observe obvious differences in SYP staining quality (puncta density and mean intensity) between cleared and non-cleared sections (data not shown). These findings from the perfusion-fixed cases could not be directly compared to the immersion-fixed cases, inasmuch as we co-stained pre- and postsynaptic proteins, and the exclusively immersion-fixed brains displayed strong autofluorescence for shorter wavelengths. Related thereto, DAPI counterstaining was only successful in the perfusion-fixed cases.

### 3.2 Increased specificity of immunostaining after CLARITY

Besides the increased sensitivity for CAMKIIA labeling after CLARITY, we observed differences in target specificity for a microtubule-associated protein 2 (MAP2) antibody, a prominent neuronal marker. MAP2 immunofluorescence on non-cleared sections revealed strong superficial labeling of undefinable, partly globular structures (especially close to section surface) not corresponding to somata or dendrites, as shown in Figure 2c (perfusion-fixed) and Figure 2d (immersion-fixed). When all six study cases were viewed in a row, the MAP2 antibody provided reliable labeling of its target structures in cleared sections independent of their fixation method and total fixation time, whereas sections without CLARITY-pretreatment showed variable and inconsistent outcomes with sometimes unspecific artifact-rich or even no specific labeling (Figure 2-figure supplement 1d).

Finally, we tested CLARITY as a substitute for historical retrieval agents, e.g., detection of tyrosine hydroxylase (TH) in neurons and processes of formalin-fixed archival human tissue, which was successful after formic acid pretreatment in (H. Braak, Thal, Matschke, Ghebremedhin, & Del Tredici, 2013). In the human neostriatum, TH is enriched in the matrix and has been shown to clearly delineate matrix/striosome organization (Morigaki & Goto, 2016). In a cross-section from the caudate nucleus, TH staining showed the described mosaic pattern for cleared sections (Figure 2c), and an inset shown in higher magnification unveiled neurites, most likely representing dopaminergic axons from nigral neurons (Morigaki & Goto, 2016), as the underlying structures. In the corresponding non-cleared section, only autofluorescent lipofuscin pigment was visible, and the tilescan (large overview scan) did not allow for clear distinction between matrix/striosome compartments.

### 3.3 3D visualization of AD-related markers and concomitant synaptic decline with CLARITY

We also tested a panel of 30 antibodies for cells and subcellular compartments as well as pathology-associated markers in combination with CLARITY to provide a comprehensive toolbox intended to complement future studies on human tissue (see also 3.4). We used only perfusion-fixed tissue from the gross anatomy course. Only antibodies that worked in at least two different cases are listed in Table 2.

First, we aimed at assessing the performance of antibodies used for neuropathological staging on cleared sections. Neuropathological AD staging is based on the regional distribution and amount of DAB-stained intraneuronal neurofibrillary tangles (NFTs) and extraneuronal amyloid-β (Aβ) deposits (H. Braak et al., 2006; H. Braak & Braak, 1991).

Since tau aggregates not only occur as NFTs, neuropil threads (NTs), and neuritic components of neuritic plaques (NPs) (Figure 3a, left image), we also investigated whether other well-characterized abnormal tau aggregates are preserved after CLARITY. AT8-positive, partially soluble, material in axons of the perforant path constitutes a special form of pretangles, referred to as axonal “non-argyrophilic abnormal tau” (H. Braak & Del Tredici, 2015). We investigated the white matter underneath the entorhinal cortex of a NFT stage III and NFT stage V case and found AT8-positive axons in cleared sections (Figure 3a, second image). In combination with Nissl staining, ghost tangles could be identified in cleared tissue based on the missing nucleus. Nissl counterstaining after CLARITY was successfully performed on cleared sections from immersion- and perfusion-fixed tissue (Figure 3a, third and fourth image).

**Figure 3.**
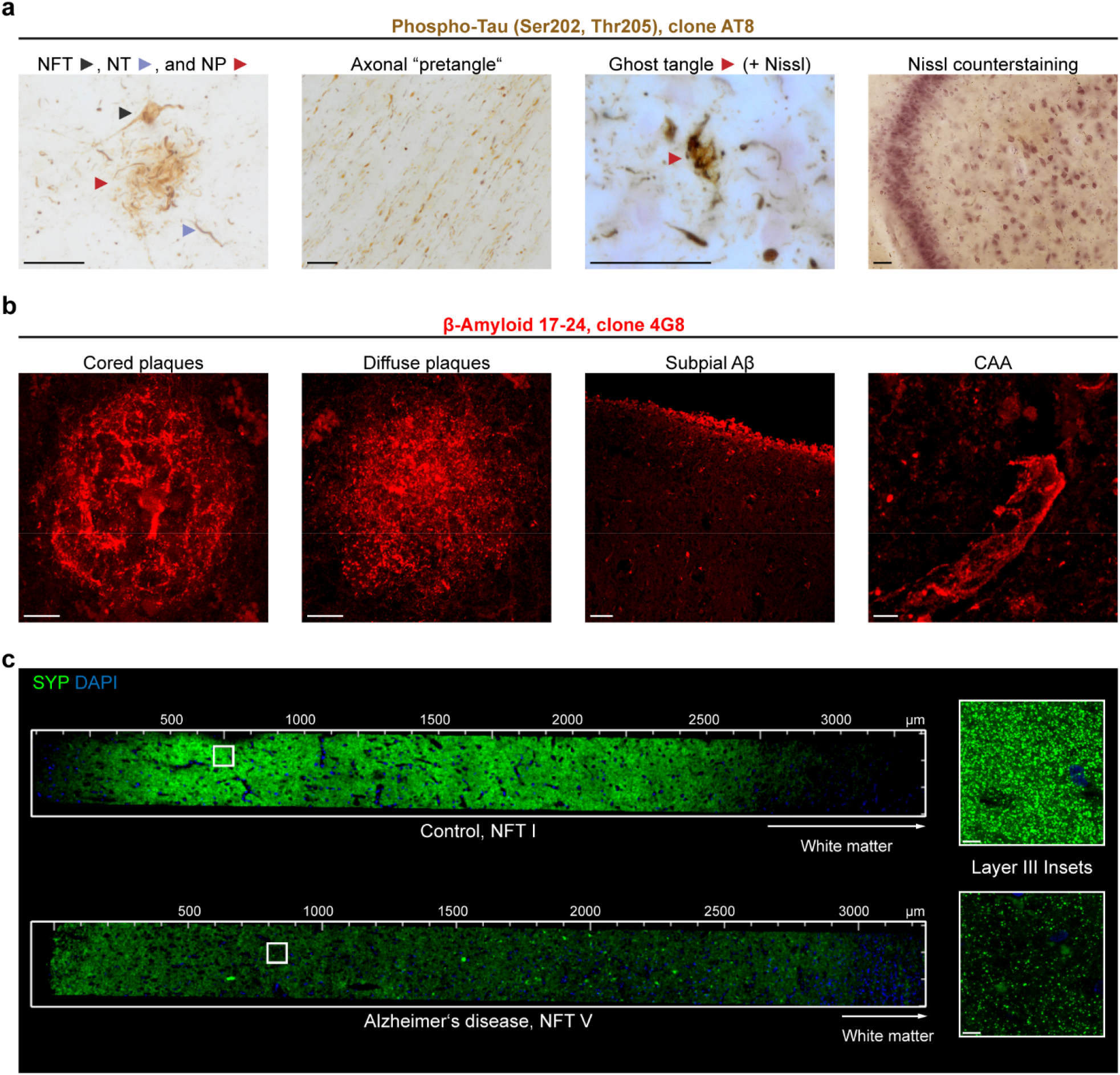
Hallmarks of Alzheimer’s disease and synaptic decline throughout the six cortical layers after CLARITY. **(a)** 100 µm-thick PEG sections (tissue bank, NFT stage V, Aβ phase 4, PD stage 3) were hydrated in 70% ethanol after cutting and washed in water before starting with the CLARITY protocol. Following the clearing step, traditional DAB-based immunolabeling was performed against phospho-tau (clone AT8). Different kinds of tau aggregates could be identified. Ghost tangles required counterstaining for Nissl substance with Darrow red. All images were acquired with a BX61 microscope (Olympus Optical) using extended focal imaging (EFI). On the far right, a cleared hippocampal section from a perfusion-fixed case is shown after AT8 staining (DAB) and Nissl counterstaining. The 2D image was acquired with a Biorevo BZ-X810 microscope. NFT = neurofibrillary tangle, NT = neuropil thread, NP = neuritic plaque. Scale bars, 50 µm. **(b)** Different types of Aβ pathology were detected following immunolabeling (clone 4G8) after CLARITY (perfusion-fixed case). CAA = cerebral amyloid angiopathy. Scale bars, *from left to right*, 10 µm, 10 µm, 50 µm, 5 µm. Except the third image from left, all images represent MIPs of z-stacks. **(c)** Staining of comparable cortex sections (superior frontal gyrus, Brodmann area 9) with synaptophysin 1 (SYP, G2 in Table 2) reveals the drastic decline in synaptic density between a control case (case p2, NFT stage I) and a clinically diagnosed Alzheimer’s case (NFT stage V). Both sections were derived from perfusion-fixed cases. The insets to the right show a magnification of the most heavily affected layer III. Scale bars, 8 µm.

The second hallmark of AD, Aβ plaques, could be identified after CLARITY in various reported morphologies (Thal et al., 2000), e.g., cored *versus* diffuse plaques, subpial Aβ, and Aβ deposits along vessels (i.e., cerebral amyloid angiopathy, CAA) (Figure 3b).

Because AD has also been described as “synaptic failure” (Selkoe, 2002), we combined the power of coherent 3D imaging and imaged all six cortical layers in the superior frontal gyrus from a control patient (case p2, NFT stage I) and an AD patient (perfusion-fixed, NFT stage V). We observed drastically reduced SYP puncta in the AD patient, which is emphasized by higher-magnified insets (Figure 3c) of the most heavily affected layer, layer III of the cerebral cortex (Scheff et al., 1990).

Some of the here presented antibodies are shown in **Videos 1-3**.

### 3.4 Antibody toolbox for human brain interrogation with CLARITY

A schematic overview on the organization of neuronal cells, their compartments, and glial cells in the central nervous system is provided in Figure 4a with examples presented in Figure 4b and c. Stainings of human archival brain tissue with the remaining antibodies from Table 2 are shown in Figure 4-figure supplement 1, followed by co-stainings and region-specific stainings in Figure 4-figure supplement 2. Tilescans of cortical and subcortical structures are shown in Figure 4-figure supplements 3 and 4. All antibodies showed expectable structures in at least two different brains.

**Figure 4.**
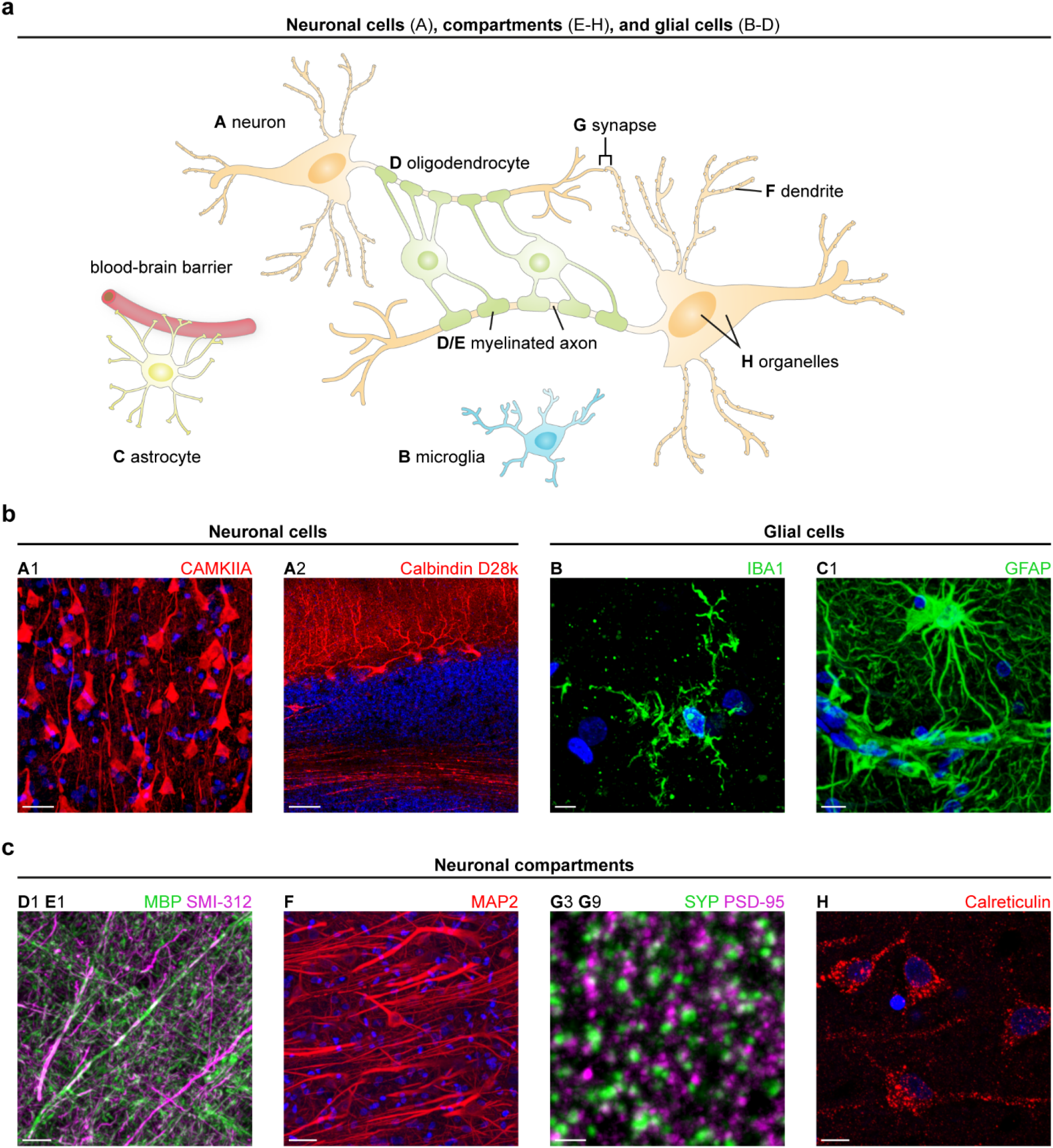
Antibodies for specific labeling of structures in the human brain. **(a)** Schematic drawing showing the interaction between neurons (A) and glial cells, e.g., microglia (B), astrocytes (C), and oligodendrocytes (D) in the central nervous system. Neuronal compartments are highlighted (E-H). **(b) (left)** Subtypes of neurons are displayed using antibodies against CAMKIIA (pyramidal cells, cerebral cortex) and calbindin D28k (Purkinje cells, cerebellum). Scale bars, 30 µm (A1), 100 µm (A2); **(right)** Antibodies against IBA1 (case p2) and GFAP label morphologically distinct cell groups. Scale bars, 5 µm (B), 10 µm (C1). **(c)** Exemplary stainings of neuronal processes (axons enveloped with myelin sheaths, dendrites), synapses (pre- and postsynapse), and organelles (endoplasmic reticulum). Scale bars, *from left to right*, 15 µm (D1 E1), 30 µm (F), 2 µm (G), 10 µm (H). DAPI counterstaining is depicted in blue. If not otherwise indicated, immunostaining was performed on frontal cortex sections from perfusion-fixed cases (case p2 for D1 E1, case p1 for F-H). Except A2, all images represent maximum intensity projections (MIPs) of z-stacks. The upper-case letter at the top of each image refers to the entities outlined in panel (a) and the combination of letter and number allows for identification of the applied antibody listed in Table 2. See also Figure 4-figure supplements 1-4.

#### Detection of distinct neuronal cells in cleared human sections

First, we set out to identify the different types of neurons in their specific brain region. As described above, the CAMKIIA antibody detected excitatory pyramidal cells in the cerebral cortex (A1 in Figure 4b). Strikingly, CAMKIIA is expressed in inhibitory Purkinje cells of the cerebellum, where somata, dendrites, and axons of Purkinje cells were labeled (Figure 4-figure supplement 3a), as reported previously for rat cerebellum (Ichikawa, Sekihara, Ohsako, Hirata, & Yamauchi, 1992). Two different antibodies against calbindin D28k successfully stained somata, dendrites, and axons of Purkinje cells, as described elsewhere (Flace et al., 2014) (A2 in Figure 4b, A3 in Figure 4-figure supplement 1a). From the vast group of interneurons, we exemplarily tested an antibody against the rather seldom used epitope, corticotropin-releasing hormone (CRH, A4 in Figure 4-figure supplement 1a) (Chen, Andres, Frotscher, & Baram, 2012) and detected interneurons in the Ammon’s horn of the human hippocampus. The TH antibody (see 3.2) was also applied in the substantia nigra (data not shown) and locus coeruleus (A5 in Figure 4-figure supplement 1a), where it highlighted the catecholaminergic principal neurons and their cellular processes.

#### Detection of glial cells in cleared human sections

Glial cells, including microglia, astrocytes, and oligodendrocytes, constitute a second large cell group in the CNS. Ependymal cells were not studied in detail here. As an alternative to the frequently applied ionized calcium-binding adapter molecule 1 (IBA1) antibody from Wako, we identified another IBA1 antibody that unveiled specific labeling of microglia in human cerebral cortex tissue, as shown in Figure 4b, B, and in a co-staining with astrocyte marker glial fibrillary acidic protein (GFAP) (B C2 in Figure 4-figure supplement 2a). Astrocytes and their enormous fiber meshwork could be traced in great detail and throughout large z-stack volumes arguing for a good preservation of the 3D organization as hypothesized previously (Lai et al., 2018).

In addition to cell somata (C1 in Figure 4b, C2 in Figure 4-figure supplement 1b), both tested GFAP antibodies impressively unveiled the contribution of the astrocytic foot processes to the blood-brain barrier (asterisks in the tilescan of Figure 4-figure supplement 4a, the white box is shown as higher-magnified image in Figure 4b, C1). The second boundary layer formed by astrocytes, the glial limiting membrane (glia limitans), located between the pia mater and the cerebral cortex, is shown in Figure 4-figure supplement 1b, C2. In a tilescan acquisition spanning 848 µm x 1.1 mm x 56 µm (in x, y, and z, respectively) of a human cerebellar section, GFAP unveiled an intricate meshwork arising from Bergmann glia with cell bodies in the Purkinje cell layer and long radial cellular protrusions in the molecular layer (Figure 4-figure supplement 4b). Whereas GFAP is expressed more broadly, S100 calcium-binding protein B (S100B)-co-expression is restricted to more mature cells with no residual stem cell characteristics (Raponi et al., 2007). Accordingly, both antibodies showed only partial overlap in a co-staining (C2 C3 in Figure 4-figure supplement 2a). In the CNS, oligodendrocytes form the myelin sheath around axons, and this could be clearly visualized in a co-staining with a pan-axonal marker (SMI-312, E1 in Figure 4c) and an antibody directed against myelin basic protein (MBP, D1). Two additional MBP antibodies were successfully applied in human tissue (D2 & D3 in Figure 4-figure supplement 1b).

Two of the here presented antibodies (GFAP and MBP) are shown in **Videos 4-5**.

#### Neuronal processes & organelles in cleared human sections

Besides the mere staining of axons (SMI-312, see Figure 4c, E1), we could visualize nodes of Ranvier with an antibody directed against a protein of the paranodal region (Figure 4-figure supplement 1c, arrows in E2). Dendrites could be traced using a MAP2 antibody (F in Figure 4c; **Video 6**) as outlined above. We aimed to complement the list of markers for intracellular organelles, as did others before, e.g., for mitochondria (Phillips et al., 2016). For detection of nuclei, DAPI counterstaining was applied in all cases. The endoplasmic reticulum could be represented with an antibody directed against calreticulin (H in Figure 4c).

Since large overview scans (tilescans) provide best insights into the cyto- and myeloarchitectonics of the cerebral cortex, we exploited this technique to obtain xyz scans extending from the outer cortical border to the beginning of the underlying white matter. By CAMKIIA-staining, layer-characteristic neuronal morphology could be highlighted on an epitope-specific level next to a sketch based on a historical drawing published by the anatomist Henry Gray (Gray & Lewis, 1918) (Figure 5a). The dendritic tufts in the molecular layer were clearly visible as well as the increasing size of pyramidal neurons from layer III to layer V. Epitope-based distinction of cortical layers represents a valuable tool, which can be combined in a co-staining with a synaptic marker to specify synaptic decline in AD. The sketch showing the myeloarchitectonics is based on a non-immuno-based myelin stain found in anatomy text books (Lüllmann-Rauch, 2009). The applied MBP antibody correlated well with the sketch, showing only little signal in the upper layers, whereas the fiber bundles in the lower layers were strongly highlighted. The transverse arrangement of fibers around layer V was reminiscent of the inner band of Baillarger. Since the cortex section (BA9) was not derived from primary sensory areas, we did not expect a clear outer band of Baillarger. The missing structural detail in the upper layers was probably due to the fact that the staining was acquired as 2D scan.

**Figure 5.**
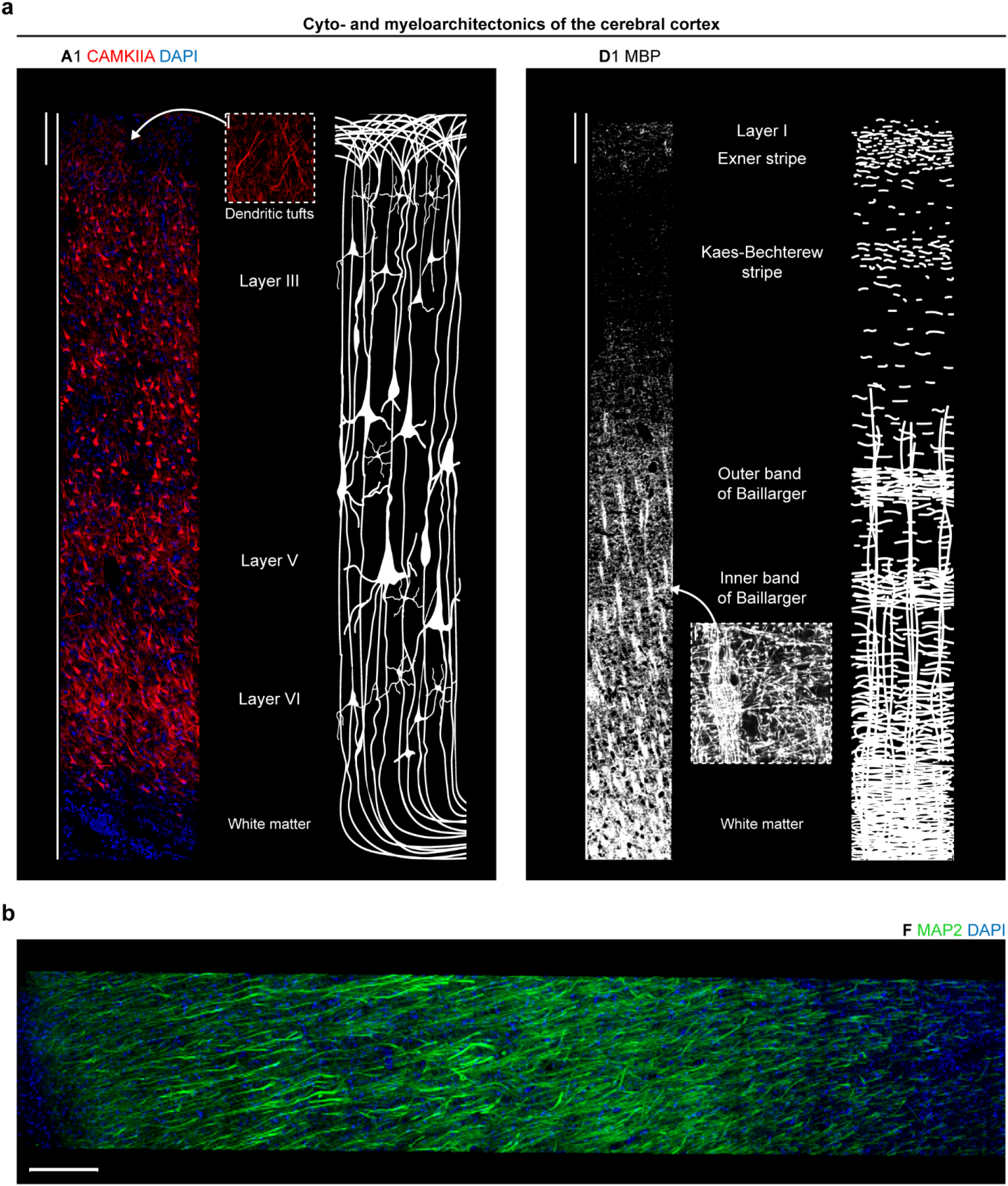
Cyto- and myeloarchitectonics in the human cerebral cortex after CLARITY. **(a) (left)** Cytoarchitectonics by 3D CAMKIIA imaging (MIP of stack with z = 25.5 µm) in the frontal cortex (perfusion-fixed) opposed to a schematic drawing based on Henry Gray. The inset (scale bar, 20 µm) shows the dendritic tufts of layer I. **(right)** Myeloarchitectonics as revealed by 2D MBP staining in the frontal cortex (case p1) opposed to a schematic drawing showing a non-immuno-based myelin stain found in anatomy textbooks. The inset (scale bar, 10 µm) shows the horizontal and transverse arrangement of fibers. Scale bars, 200 µm. **(b)** MAP2 immunostaining in the superior frontal gyrus of case p1, MIP of stack with z = 25.5 µm. Scale bar, 200 µm.

In addition to cell and myelin organization, we captured the network of MAP2-positive dendrites and cells in a tilescan of the frontal cortex (Figure 5b). In a z = 63 µm tilescan showing the molecular layer of the cerebellum, MAP2 allowed for a clear rendering of dendrites from Purkinje cells and somata of interneurons (Figure 4-figure supplement 3b). Remarkably, the CAMKIIA and MAP2 antibody can be also applied for precise determination of the gray/white matter border because the immunoreactivity of both antibodies remarkably dropped where the approximate beginning of the white matter was marked with a pen on the slide, leaving only minor signal from interstitial neurons (see (Suarez-Sola et al., 2009) for a review). This finding underlines the target-specificity of both antibodies.

#### Successful detection of synaptic subtypes in human brain sections

Since synaptic loss has been described as a molecular correlate of cognitive decline in neurodegenerative diseases like AD (DeKosky & Scheff, 1990; Terry et al., 1991) and ALS (Henstridge et al., 2018), excitatory and inhibitory synapse markers were tested, which is demanding in long-term formalin-fixed tissue. For overall staining of all synapses, the presynaptic protein SYP has been a prominent target as outlined above. Both SYP antibodies tested here showed the typical punctate pattern and a strong overlap in a co-staining, as seen in Figure 4-figure supplement 2a, G2 G3. The partial overlap can be explained by targeting different epitopes of SYP (i.e., different isoforms). In the same category of antibodies, we successfully applied a synapsin 1/2 antibody (G1 in Figure 4-figure supplement 1c).

To highlight the presynaptic site of exclusively inhibitory synapses, a cleared section from the putamen was stained for vesicular gamma-aminobutyric acid (GABA) transporter (VGAT). In line with a previous study (Dunn et al., 2019), the punctate pattern could be also observed in the somata of medium spiny neurons and/or GABAergic interneurons (G4 in Figure 4-figure supplement 1c). Robust results were obtained with a VGLUT1 antibody for detection of excitatory synapses. Since VGLUT1 has been described as being enriched in the human putamen but absent in the pallidum (Vigneault et al., 2015), we subjected the antibody to a specificity test in both regions and found consistent expression patterns (Figure 4-figure supplement 2d, II G5 putamen and III G5 pallidum). Moreover, in a tilescan covering the complete cortex (superior frontal gyrus), VGLUT1 (Figure 4-figure supplement 2e) and SYP (Figure 3c) were nearly absent in the underlying white matter.

We next tested several postsynaptic antibodies on cleared sections, a challenging endeavor in formalin-fixed archival tissue. An anti-gephyrin antibody was applied in the pallidum (G6 in Figure 4-figure supplement 1c). To our knowledge, GLUA2 has not yet been routinely analyzed in immunolabeling studies on human brain tissue. The antibody in Table 2 not only showed a good co-localization with a presynaptic marker (G7 G2 in Figure 4-figure supplement 2b) but also unveiled detailed expected substructure in super-resolution microscopy (Figure 7, see further descriptions in 3.6). Moreover, when applied in western blots, it detected its target protein in commercially available human tissue lysates (Figure 4-figure supplement 2b).

From the HOMER protein family, we identified a reliably working HOMER3 antibody. Whereas HOMER1 and 2 are widely expressed in the mouse brain, murine HOMER3 expression is mainly restricted to the cerebellum and to the hippocampus (minor levels compared to cerebellum) (Shiraishi-Yamaguchi & Furuichi, 2007). In line with DAB-stained human sections (Jarius & Wildemann, 2015), Purkinje cell somata, dendrites, spines, and axons were labeled (Figure 4-figure supplement 1c, G8). The structural compartments of PSDs, the spines, were highlighted in remarkable quality along the dendrites, as seen in Figure 4-figure supplement 3b, inset G8 and **Video 7**. The PSD-95 antibody also showed good co-localization with presynaptic SYP in cleared sections (G3 G9 in Figure 4c).

Cell- and compartment specificity as well as the advantages of multiple simultaneous labeling could be demonstrated in a triple-staining of the hippocampus, a key player in neurodegenerative disorders, such as AD (A1 E1 and A1 G2 in Figure 4-figure supplement 2c). Whereas the pan-axonal marker SMI-312 traced the fibers from the alveus and highlighted outgoing mossy fibers at the hilus, the CAMKIIA antibody highlighted pyramidal cells in the Ammon’s horn and showed punctate labeling, e.g., in *stratum lucidum* and *stratum oriens*. Co-localization with the presynaptic protein SYP was especially prominent in CA3, as shown by the white overlap in Figure 4-figure supplement 2c (right-hand image, higher-magnified inset).

#### Additional targets in human brain sections

Since AD pathology can also be associated with vascular structures, e.g., CAA, we tested vessel-related markers on cleared sections. Two antibodies designed for application in human tissue gave strong signals and a penetration depth of at least 70 µm, one of which was raised against collagen IV of basal laminae, the second against α-actin of smooth muscle cells (J1 & J2 in Figure 4-figure supplement 1d; **Videos 8-9**). From a biochemical approach, a metabolism-related antibody against glyceraldehyde-3-phosphate dehydrogenase (GAPDH), an enzyme involved in glycolysis (K in Figure 4-figure supplement 1d), was also applied successfully.

### 3.5 Re-staining of fluorescently-labeled sections

Given the restricted availability of human *post mortem* brain tissue, the limited size of blocks through different brain regions, and especially analyses that require more than three to four different channels, multiple round labeling of already stained sections would be beneficial. As shown for mouse brain sections in the original CLARITY publication (Chung et al., 2013), de-staining in clearing solution and subsequent re-staining is feasible. We tested this option on perfusion-fixed tissue and stained a cortical section for pre- and postsynapses (SYP and GLUA2). We successfully performed a second round of staining with cellular antibodies (Figure 6). Most important, a control section mounted and imaged after de-staining did not show any specific signal, nor was DAPI staining detectable any longer (Figure 6). To also control for detachment of primary antibodies, the control section was subjected to the second round of staining, starting with secondary antibody incubation. Confocal microscopy did not show any signal except for re-stained nuclei.

**Figure 6.**
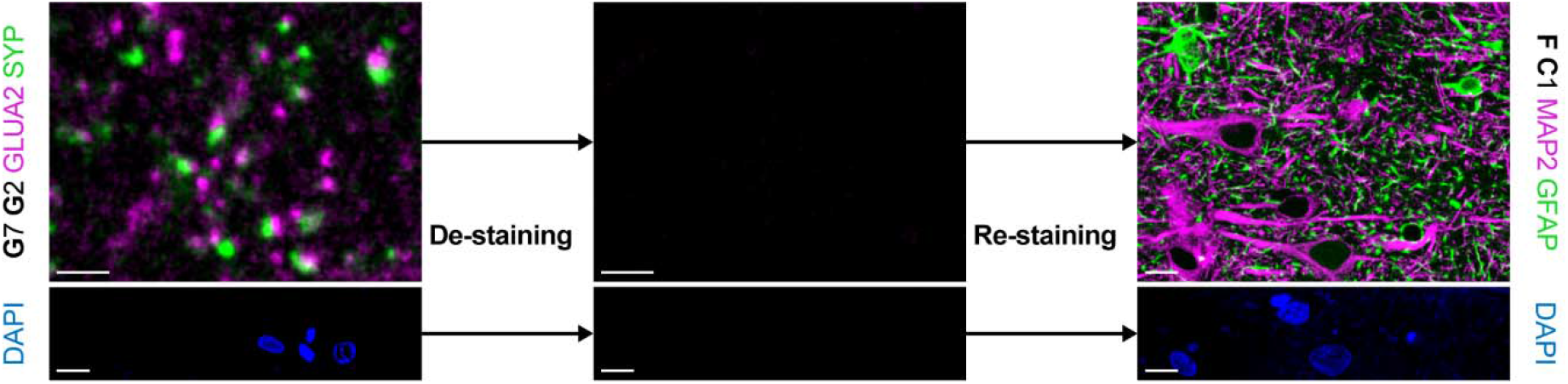
Multiple staining rounds with hCLARITY. **(top row)** Stained sections from case p1 mounted in SlowFade were de-stained in clearing solution for 24 hours at 60 °C (*left*). A de-stained section was mounted and imaged as control (*middle*). Re-staining with cellular markers (*right*). Scale bars, 2 µm (*left and middle*), 10 µm (*right*). **(bottom row)** DAPI channel of the images shown in the top row before de-staining (*left*), after de-staining (*middle*), and after re-staining (*right*). Imaging parameters were identical. Scale bars, 10 µm.

### 3.6 Super-resolution microscopy of neurons and synapses

Finally, we applied tested antibodies for super-resolution microscopy to gain insights into details that are not accessible with diffraction-limited light microscopy. To this end, we employed STED (Osseforth et al., 2014) and dSTORM microscopy (Schoen et al., 2015) with custom-built setups. First, we performed STED microscopy on GLUA2-stained sections due to the good performance of the antibody in confocal microscopy. In direct comparison to confocal images, we saw finer structures in super-resolved images (Figure 7a). While in the confocal images all synaptic puncta had homogenous shapes, the corresponding STED images unveiled the known disk shape (in side view appearing as bars (Dani, Huang, Bergan, Dulac, & Zhuang, 2010), full width at half maximum (FWHM) was 118 nm in STED vs. 594 nm in confocal) and further resolved the substructures within a single PSD. The latter could be reflected by the corresponding intensity profile, where a homogeneous intensity profile for confocal microscopy is split into two individual peaks in STED microscopy (Figure 7b).

**Figure 7.**
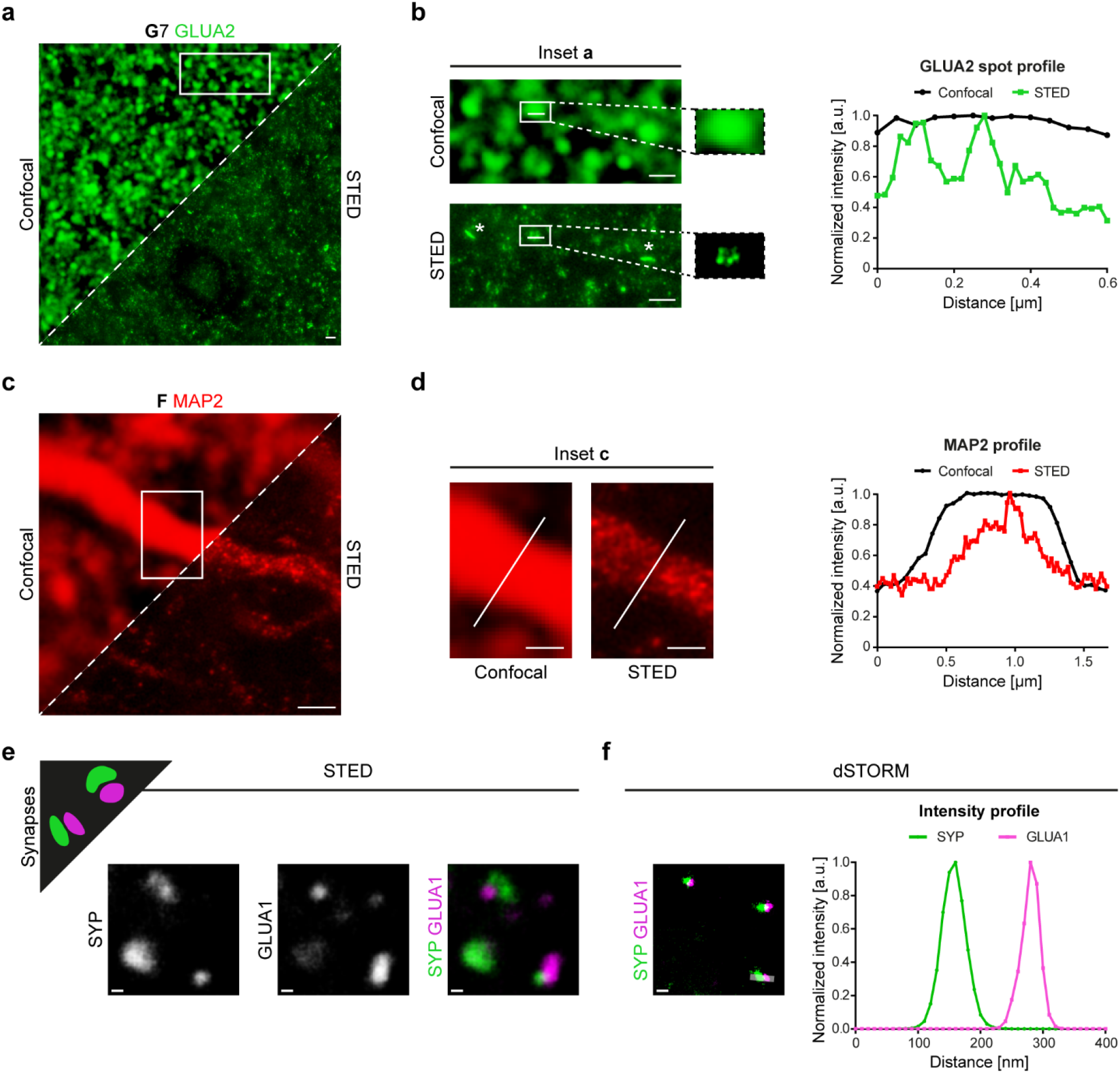
Super-resolution microscopy with STED and dSTORM imaging of synaptic and neuronal markers on human brain tissue after CLARITY. **(a)** Synaptic GLUA2 in the cerebral cortex acquired in confocal (*upper left*) and STED mode (*lower right*). Scale bar, 1 µm. **(b)** Insets (boxed area from (a)) provided in higher magnification unveil postsynaptic membrane morphologies like disk shape as seen in a side-view on a synapse (*asterisks, left column*) and the detailed localization of GLUA2 within a single synaptic spot as seen in a top view on a synapse (*right column*). A line intensity profile (*right part* of (b)) across the spot surface demonstrates the intensity fluctuation along the spot (*white line in the boxed areas* of (b)). Mean intensities (gray values) were normalized on the highest value. a.u. = arbitrary unit. Scale bar, 1 µm. **(c)** MAP2 staining in a cerebral cortex section. STED and confocal modes of the same neuron are shown in comparison as in (a). Scale bar, 1 µm. **(d)** Insets (boxed area from (c)) provided in higher magnification show structural details in the STED mode. The intensity profile across the white lines in the images is plotted on the right. Scale bar, 0.5 µm. Sections for MAP2 and GLUA2 staining were derived from the superior frontal gyrus of perfusion-fixed cases. In (a)-(d), pixel size for confocal images is 50 nm. **(e)** Schematic drawing showing expected precision imaging of synaptic compartments in super-resolution images (*upper left*). SYP/GLUA1 co-staining in STED microscopy. The channels are shown individually (*left and middle*) and as overlay (*right*). Here, sections were incubated in hydrogel for two days, total clearing time was three days. Scale bars, 200 nm. **(f)** SYP/GLUA1 co-staining in dSTORM microscopy (*left*). An intensity profile across a synapse (rectangle in the left image, width of ten pixels) is shown on the right. Sections were incubated in hydrogel for one day, total clearing time was two days. Scale bar, 200 nm. All sections in (e)-(f) were derived from perfusion-fixed cases. See also source data file and Figure 7-figure supplement 1.

Second, while the MAP2 antibody showed a rather continuous signal in confocal images (FWHM of entire dendrite = 1.21 µm), the corresponding STED image unveiled structural details with a punctate pattern of the signal (FWHM = 521 nm) (Figure 7c), which was well comparable to STED images obtained from cultured neurons with the same antibody (Leschik et al., 2019). The intensity profile across a dendrite showed a narrower peak (on a broader background), highlighting the superior resolution of STED (Figure 7d).

Third, to trace synaptic and neuronal structures in 3D, a series of images was acquired from a CAMKIIA-labeled section. Similar to MAP2 labeling, the detailed organization of CAMKIIA-positive structures and clusters in dendrites could be studied in the maximum intensity projection image (Figure 7-figure supplement 1 and **Video 10**). Finally, in a proof-of-principle staining, we tested the compatibility of dSTORM imaging with a pre- and postsynaptic staining after CLARITY and opposed it to a STED image (Figure 7e) labeled for the same compartments (SYP antibody for presynapses, GLUA1 antibody for postsynapses). Synapses were presented in high detail and we obtained better localization precision and hence increased resolution between two closely localized protein clusters with dSTORM images (Figure 7f).

## 4. Discussion

### 4.1 Major advantages of hCLARITY

The main findings of this study were, first, that only with the additional application of CLARITY prior to IHC, immunostaining with some markers (MAP2, TH) was possible (i.e., more specific). With other markers (CAMKIIA, GLUA2), staining became more intense, not only in depth, as was known and expected with CLARITY, but already at the surface (i.e., more sensitive). Second, we offer a comprehensive set of 30 different well-working antibodies. Furthermore, sections can be completely de-stained (including DAPI) and re-stained. Third, super-resolution microscopy with two different techniques was possible, for the first time also at the level of synapses in the human brain. Furthermore, we could show that hCLARITY can not only be combined with modern fluorescence-based microscopy techniques but also with chromogen-based staining and Nissl staining. Perfusion-fixed tissue was derived from body donors of the gross anatomy course, which is a valuable source of *post mortem* tissue, for some aspects superior to solely immersion-fixed tissue (less autofluorescence in the shorter wavelength spectrum). The method is particularly attractive because it is inexpensive and easy to perform.

It should be emphasized that no loss of detectability of various AD tau and Aβ protein aggregates was observed with the CLARITY protocol applied here (hCLARITY). Already in the initial publication of the method, CLARITY was shown to be applicable to longer fixed human brain tissue for IHC staining (Chung et al., 2013). Applicability to sections has also been previously described (Poguzhelskaya, Artamonov, Bolshakova, Vlasova, & Bezprozvanny, 2014). Taken together, in addition to the known increase in antibody penetration, we have identified further benefits of the CLARITY method in the sense of antigen retrieval by enhancing or enabling accessibility of epitopes in long-term fixed human brain tissue. Therefore, we propose that hCLARITY be applied prior to IHC.

We have presented 30 usable primary antibodies directed against different cell types, cellular structures, and neurodegenerative pathologies. Each of these antibodies proved to be robust, passing quality control only when functioning in brains of at least two individuals. For markers with no clearly identifiable localization (e.g., synaptic markers), the quality threshold was reached only when co-localizing with another synaptic marker. We compared the obtained expression patterns with available literature and found a good match, indicating the compatibility of hCLARITY with these antibodies. This is important, given the fact that staining of IBA1 was not possible with a clearing method which included OPTIClear (Lai et al., 2018), whereas in the present study using hCLARITY the staining was successful. This panel represents one of the most comprehensive ‘antibody toolboxes’ for immunofluorescence microscopy published to date for human brain research. The primary antibodies described here can be used for co-staining as an initial specificity test for antibodies applied for the first time on human brain tissue and also as a test for tissue quality. Previous studies applying methods, such as array tomography or CLARITY, and additional selected clearing techniques for immunofluorescence staining of the human brain, have usually presented significantly fewer antibodies, and the focus of these publications has mostly been on other aspects (20 primary antibodies in (Kay et al., 2013); 4 with CLARITY in (Ando et al., 2014); 4 with CLARITY in (A. K. Liu et al., 2016); 2 with CLARITY from (Costantini et al., 2015); 10 with CLARITY from (Phillips et al., 2016); 2 apart from several amyloid and tau antibodies in (Liebmann et al., 2016); 4 in (A. K. L. Liu, Lai, Chang, & Gentleman, 2017); 9 with CLARITY in (Morawski et al., 2018)).

In this regard, we consider the work of Ando et *al.* (Ando et al., 2014) as a forerunner of our own work; this group pioneered the use of CLARITY for 3D imaging of AD-associated neuropathological changes in tissue, hinting at its true potential. The authors were already able to show that CLARITY appeared to be partially superior to other clearing techniques for purposes of visualizing such changes (Ando et al., 2014). In previous studies, e.g., (Phillips et al., 2016), significantly longer antibody incubation times were used, but our focus was not on the depth of penetration and reduction of light scattering but, rather, on the quality of the IHC. In contrast to Ando et *al.* (Ando et al., 2014), we mostly used the commercial mounting reagent ProLong Gold antifade (refractive index (RI) after curing = 1.46) for cleared sections. In comparison to self-made RIMS, consistent formulation is guaranteed by the vendor. Further, we frequently observed crystal formation days after mounting in RIMS, whereas ProLong Gold antifade mountant enables long-term storage with concurrent preservation of fluorescence. Our clearing times were similar to those used in the study by Magliaro et *al.* who found that significantly longer detergent treatments can lead to antigen loss (Magliaro et al., 2016).

An additional discovery that emerged from the present work is the application of the hCLARITY procedure for manifold IHC and microscopy techniques, which is especially valuable in light of the restricted access to human *post mortem* tissue. Complete de-staining is already possible within one day, but has not always been reproducible with human tissue (compare (Phillips et al., 2016) and (A. K. Liu et al., 2016)). Here, re-staining with the use of a completely different antibody combination (synaptic to cellular) functioned without any background or remaining residual staining from the first staining. The advantages for studying neuropathological changes are obvious; for example, neurodegenerative protein deposition and changes in synaptic integrity (first staining) and cells (second staining) can be studied on the same tissue section. To our knowledge, ours is also the first study to show that a DAPI stain is also completely de-stainable; thus, another channel can be used in a subsequent round of labeling or, if necessary, the tissue section can be re-stained with DAPI.

Besides diffraction-limited confocal microscopy, we showed human brain synapses in sub-diffraction resolution in the super-resolution microscopy techniques STED and dSTORM. Owing to the higher spatial resolution, a higher labeling density and a lower background signal is required in super-resolution optical microscopy as compared to classical fluorescence microscopy approaches. Therefore, the requirements with respect to clearing conditions of tissue are much more stringent, thus providing an important quality test of hCLARITY. Protein aggregates in neurodegenerative diseases, e.g., tau deposition in human AD samples, have already been shown in a previous work using STED (Benda, Aitken, Davies, Whan, & Goldsbury, 2016). Here, to our knowledge for the first time, we could show a full chemical synapse with pre- and postsynaptic markers in a human *post mortem* brain with super-resolution microscopy, as has been previously demonstrated only in mouse brain (Dani et al., 2010) or in rat hippocampal neurons (Schoen et al., 2015) with dSTORM. Furthermore, in our hands, STED imaging with GLUA2 allowed unprecedented insights into the molecular organization of human postsynapses in the cerebral cortex, with a FWHM comparable to data obtained in mouse brain (disk shape: 118 nm here vs. 90 nm in (Masch et al., 2018)). The distinct arrangement of GLUA2 puncta within one PSD might reflect the concept of nanodomains (Hruska, Henderson, Le Marchand, Jafri, & Dalva, 2018; Nair et al., 2013).

The implications of these findings for neurological disease research are manifold, e.g., the still unresolved questions of pre-or postsynaptic localization of proteins that may be altered in neurodegenerative diseases, such as amyloid precursor protein (APP) (Schubert et al., 1991; Westmark, 2013), or huntingtin (Gutekunst et al., 1995), or the study of the mechanisms of network anterograde transsynaptic transmission of misfolded proteins (H. Braak & Del Tredici, 2016).

### 4.2 Beneficial mechanisms of hCLARITY on long-term-fixed human brain tissue

Presumably, the CLARITY method has some advantageous chemical properties when using formalin-fixed archival brain tissue. In particular, CLARITY has been shown to enhance the penetration depth of immunostaining (Chung et al., 2013), including human brain tissue (Morawski et al., 2018). On the one hand, this may be due to the increase in transparency; on the other to the better penetrability of the tissue for antibodies after removal of the biolipids. When considering the benefits of CLARITY, rather too little attention was paid to the fact that SDS treatment appears to increase the immunogenicity of antigens, presumably through denaturing properties (Robinson & Vandre, 2001; Syrbu & Cohen, 2011). Importantly, descriptions from preliminary studies have indicated that most of the tested antibodies showed either improved or unaltered labeling after SDS treatment (Brown et al., 1996; Robinson & Vandre, 2001; Salameh, Nouel, Flores, & Hoops, 2018). While it has already been speculated about a retrieval effect of SDS in the framework of CLARITY (A. K. Liu et al., 2016), we demonstrated that hCLARITY significantly increases the sensitivity, accuracy (specificity), and comparability of tissues of different origin and fixation type (short/long fixation, fixation via perfusion or immersion). In line with the effects of SDS treatment alone, described by Brown et *al.* (Brown et al., 1996), we identified antibodies, that (1) did not specifically label their target proteins without hCLARITY (MAP2, TH), (2) showed stronger (more sensitive) labeling of target structures after hCLARITY (CAMKIIA, GLUA2), and (3) did not benefit from hCLARITY (SYP).

Because in laser-scanning microscopy, higher laser powers inevitably raise the proportion of tissue autofluorescence in the acquired signal (e.g., lipofuscin), more sensitive staining after hCLARITY is beneficial in this context, allowing for a decrease in laser power while keeping antibody concentrations unchanged.

#### Ideas and Speculations

The hCLARITY method has additional merits, however, inasmuch as the hydrogel stabilizes the tissue, so that modifications can be employed depending on the application, e.g., longer clearing/antibody incubation times if 3D aspects are particularly significant for the question being addressed. Multiple staining rounds with de- and re-staining are also likely to benefit from tissue stabilization. In brain tissue, there is the additional factor of its high fat content. Owing to the physicochemical removal of these lipids, the tissue probably benefits particularly strongly from the method.

We also observed that hCLARITY reduces non-specific staining of some antibodies (e.g., globular structures in MAP2 staining). It is possible that detergent treatment removes sticky particles that have been deposited. In addition, CLARITY may render individual epitopes more accessible not only by denaturation but also by mild disruption of extremely cross-linked PFA-protein networks after long fixation. The precise effect of SDS is most likely mediated on epitope level rather than on antigen level (e.g., as shown for two different caveolin-1 antibodies in (Robinson & Vandre, 2001)).

### 4.3 Limitations

We have not investigated in greater detail whether there are other beneficial properties in addition to those described here, e.g., deeper penetration upon increasing duration of clearing or the incubation times for antibodies. We also do not know yet how many times in succession de- and re-staining can be performed. The role of SDS treatment alone without prior hydrogel formation (Free of acrylamide SDS-based tissue clearing (FASTClear), (A. K. L. Liu et al., 2017)) has not been separately investigated, but two disadvantages are likely: Transparency is lower because hydrogel might also contribute to RI alignment, and tissue integrity might also suffer from prolonged SDS treatment without prior stabilization. In previous descriptions of sole SDS treatment prior to immunolabeling, much lower SDS concentrations have been used (Brown et al., 1996; Robinson & Vandre, 2001). Thus, it could be that higher SDS concentrations require chemical stabilization, e.g., via a hydrogel scaffold.

### 4.4 Future applications of hCLARITY for neuropathology research

We view hCLARITY as a significant extension of the existing spectrum of methods for research of the human brain, especially when tissue properties are not perfectly controllable. The applicability to long and differently fixed tissues will make tissue bank resources accessible to more researchers. As an example, we point in particular to the advantages of hCLARITY as an upstream method for IHC in the study of neurodegenerative diseases. Until now, findings from classical IHC were often limited to the distribution of protein deposits in various brain areas and cell compartments (H. Braak & Braak, 1991; H. Braak & Del Tredici, 2016; H. Braak et al., 2003; Brettschneider et al., 2013). By contrast, changes in structures important for neural tissue functionality (cell densities, synaptic integrity) have often been difficult to compare or have been studied using very elaborate techniques that tend to be available only in specialized laboratories. The hCLARITY method can be combined with classical histological staining (shown here with Nissl) and with DAB chromogenic immunostaining. Importantly, the use of immunofluorescence, with the additional possibility of multiple round labeling, permits the examination of several markers simultaneously (or consecutively) on the same tissue section in a quantitative and specific manner. Thus, hCLARITY can be used to study neuropathology (protein deposition) affecting functionally important structures (cells, synapses) quantitatively on one tissue section in three dimensions. The possibility of using super-resolution microscopy also expands the arsenal of methods for investigating even at the molecular level the propagation of pathologies. Recent studies with super-resolution microscopy revealed novel insights into the composition and localization of protein aggregates in AD (Querol-Vilaseca et al., 2019) and Parkinson’s disease (PD) (Colom-Cadena et al., 2017; Moors et al., 2021; Shahmoradian et al., 2019).

## 5. Conclusions

In closing, the chief advantage of hCLARITY is not only the increase of antibody and light penetration depth but also restoration of the quality of long-fixed tissue. This method and the provided information on applicable microscope systems could serve to study neuropathological phenomena in the human brain in much greater detail in the future with the here presented neuronal, non-neuronal, synaptic, and pathology markers (n = 30). The hCLARITY methodology can be performed with a basic confocal microscope at low time and with minimal material cost in any molecular biology laboratory ranging from basic research to neuropathological diagnostics. In particular, we recommend the use of perfusion-fixed brain tissue from permanent body donors, as it is often available in anatomical institutes and partially superior to immersion-fixed tissue.

## 6. Author contributions

TMB and MS designed the study. SW contributed to the details in the design and performed most of the experiments. SF and MS performed some of the experiments. DD, SW, and MS made the STED acquisitions. DD and MS conducted the dSTORM acquisitions. JM supervised all super-resolution microscopy experiments. FR and KD contributed in discussions and ideas. KD gave MS the opportunity to learn the CLARITY method in his lab, and he and his group provided expertise for application to human brain tissue. The first draft of the manuscript was written jointly by SW and MS and all authors commented on previous versions of the manuscript. All authors read and approved the final manuscript.

## 7. Data availability

This study includes no data deposited in external repositories. Data not included in the manuscript may be released upon reasonable request to the corresponding author and in compliance with the ethics vote.

## 9. Acknowledgements

We thank Ursula Pika-Hartlaub for excellent technical assistance. We also thank Sarah Mackert, Stefanie Suhm, Harun Asoglu, and Emanuel Keller for their support in the initial establishment of the CLARITY method in the laboratory. We thank Prof. Dr. Frank Kirchhoff for permission to use the Zeiss confocal microscope and Prof. Dr. Bernd Knöll for access to the Biorevo microscope. We also thank Prof. Dr. Heiko Braak and Dr. Dr. Kelly Del Tredici-Braak for fruitful discussions. Special thanks go to the body donors of the Institute for Anatomy and Cell Biology and of the tissue bank collection at Ulm University. Without their generosity these kind of studies would not be possible.

## 11. Ethics approval

This study was performed in compliance with the ethics committee guidelines of Ulm University as well as German federal and state law governing human tissue usage. Informed consent was obtained for all cases.

## 13. Figure supplements

**Figure 2-figure supplement 1.**
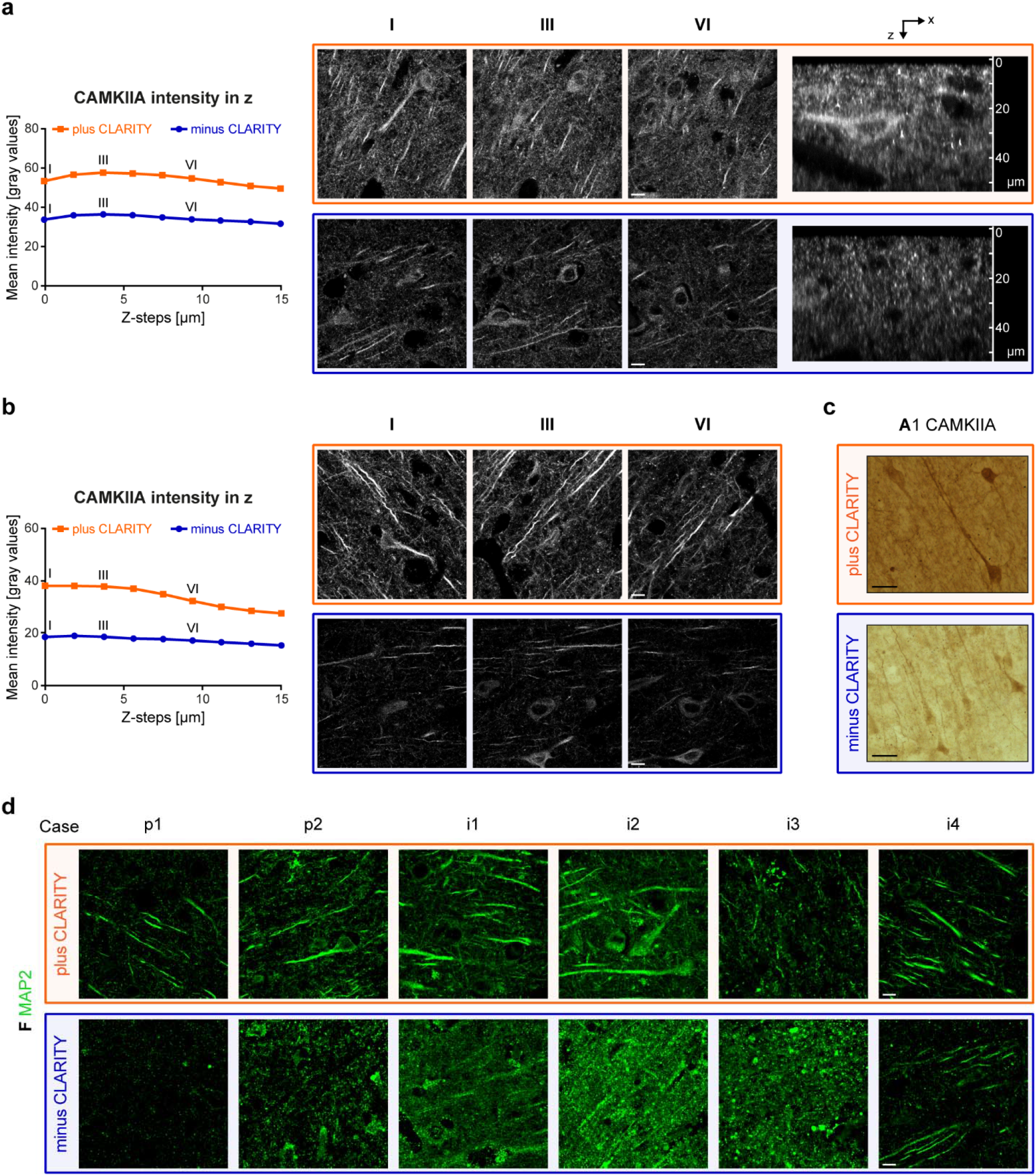
Improved antibody labeling and better comparability between different cases with CLARITY. **(a)** Cleared (orange) and non-cleared (blue) frontal cortex sections (superior frontal gyrus) from case i4 (total fixation time 18 years) were stained for CAMKIIA and imaged 15 µm deep in 1.8 µm steps using a confocal microscope. Mean intensities of each z-step were exported with LAS-X software from two stacks (cortex layer V) and the mean was plotted in dependency of the z-level. Z-level I (z = 0 µm), III (z = 3.6 µm), and VI (z = 9 µm) are provided as single images. A side view (xz scan) is shown on the far right. Scale bars, 10 µm. **(b)** Mean intensity of CAMKIIA staining on case i1 (total fixation time 5 years) was plotted in dependency of the z-level. Acquisition strategy is the same as outlined in (a). Scale bars, 10 µm. **(c)** CAMKIIA staining with DAB on cleared (orange) and non-cleared (blue) sections (case p2). Images represent full focus projections of z-stacks with identical brightness/contrast adjustments. Scale bars, 20 µm. **(d)** Comparison of MAP2 labeling on cleared (orange) and non-cleared (blue) sections from the six study cases with different fixation methods and total fixation times (see Table 1). Scale bars, 10 µm. The codes A1 and F in panels (c) and (d) refer to Figure 4. See also source data file.

**Figure 4-figure supplement 1.**
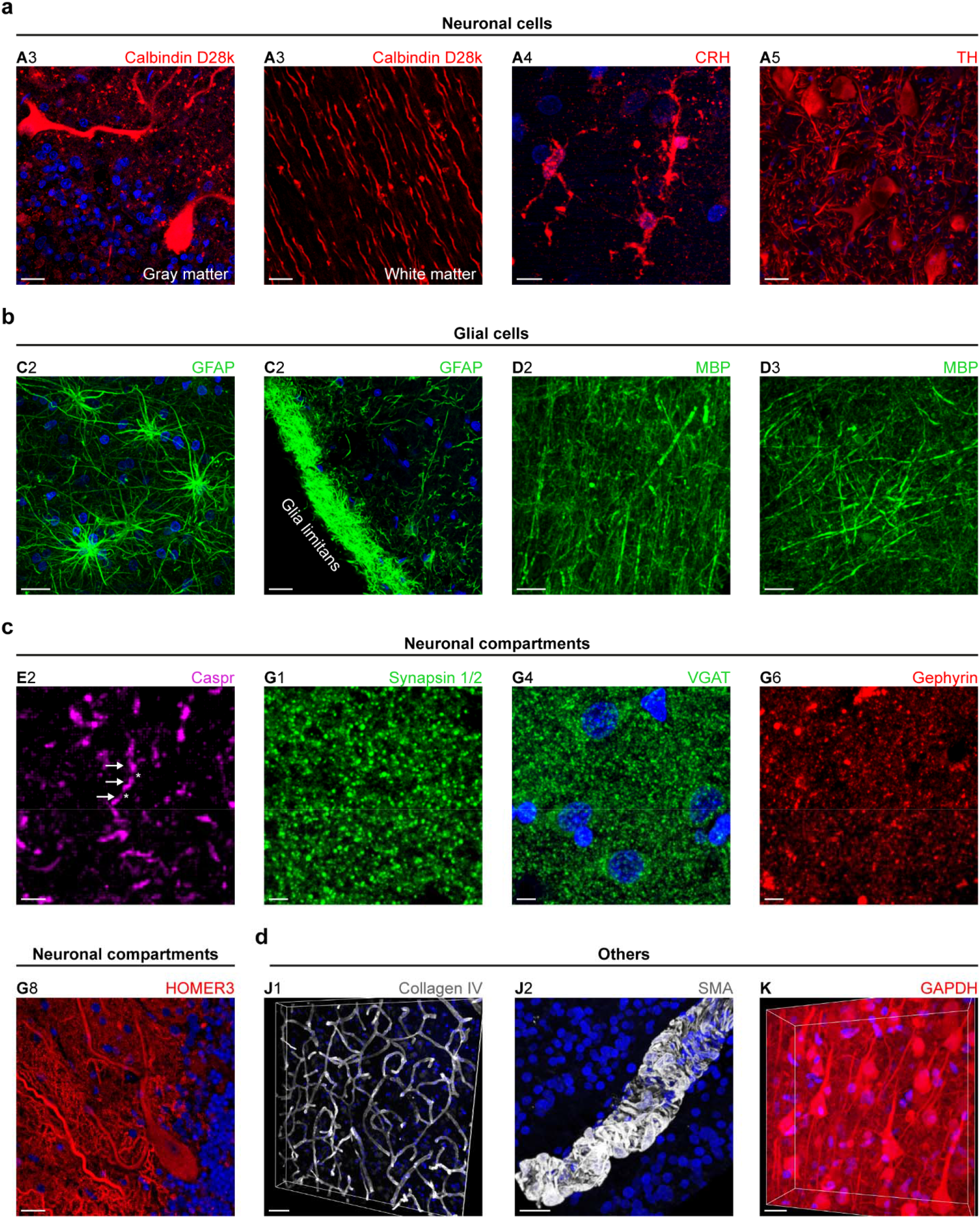
Additional CLARITY-compatible antibodies on human brain sections. **(a)** Neuronal cell markers used to detect specific neurons in cerebellum (A3), hippocampus (A4), and locus coeruleus (A5). Scale bars, 20 µm (A3), 10 µm (A4), 30 µm (A5). **(b)** Glial cell markers used on frontal cortex sections. Scale bars, 20 µm (C2), 30 µm (D2, D3). **(c)** Neuronal compartments, e.g., paranodes of axons (E2, exemplarily marked with arrows, asterisks mark nodes of Ranvier), presynapses (G1, G4), and postsynapses (G6, G8) are highlighted in frontal cortex sections (E2, G1), putamen (G4), pallidum (G6), and cerebellum (G8). Scale bars, 3 µm (E2), 5 µm (G1, G4, G6), 20 µm (G8). **(d)** Vessel-related markers (J1, J2) and GAPDH (K) in cortical sections. Scale bars, 50 µm (J1), 20 µm (J2, K). DAPI counterstaining is depicted in blue. All images except A3 (both) and C2 (right) represent MIPs of z-stacks, K is shown in perspective view. Staining is shown for perfusion-fixed cases (mostly p1 and p2, except A3, G4, and J2). The upper-case letter at the top of each image refers to the entities outlined in Figure 4a and the combination of letter and number allows for identification of the applied antibody listed in Table 2.

**Figure 4-figure supplement 2.**
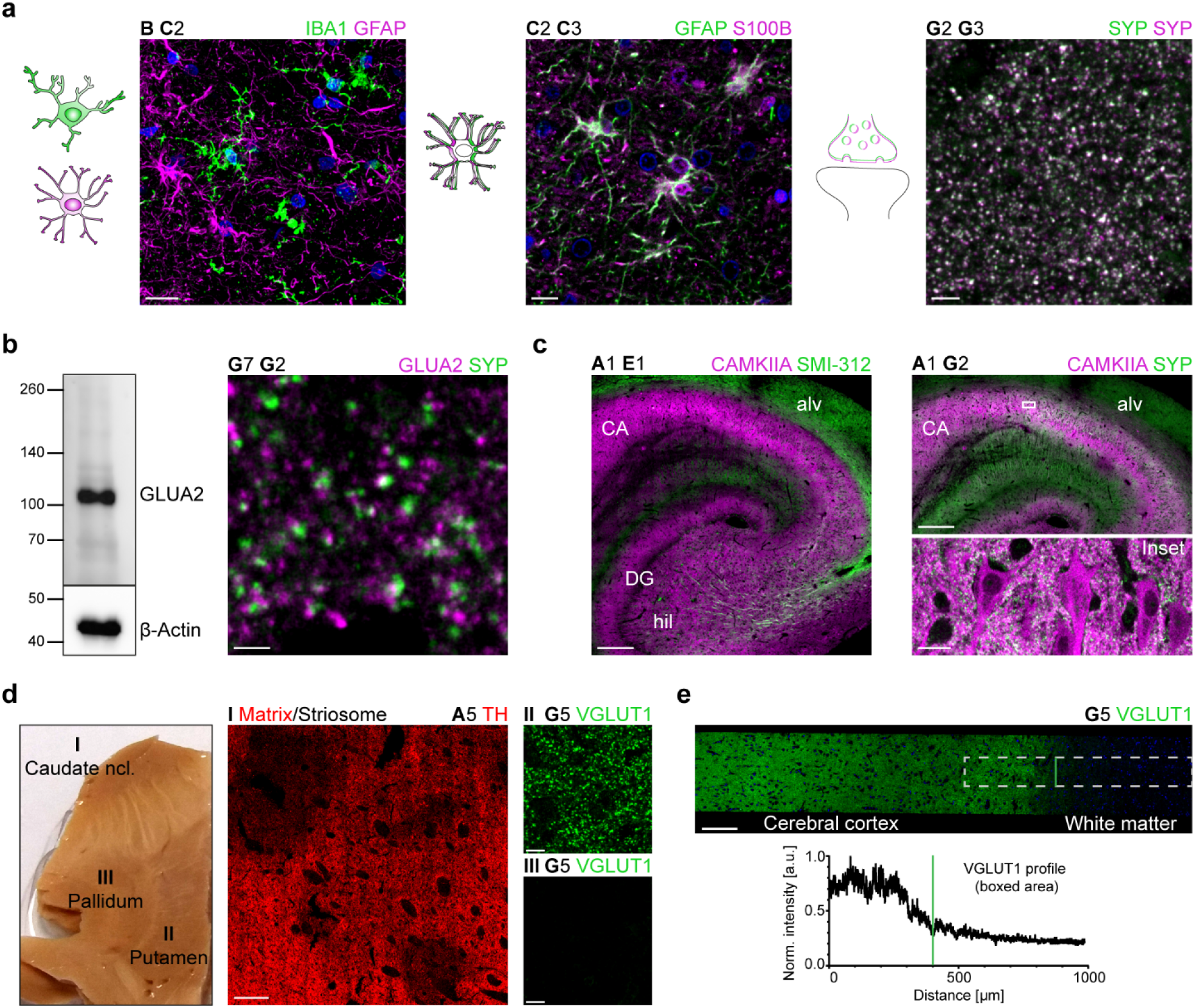
Cell, compartment, and region specificity of applied antibodies with CLARITY. **(a)** Co-staining with glial cell markers and two presynaptic markers as proof-of-specificity. DAPI counterstaining is shown in blue. Scale bars, 15 µm (*left*), 10 µm (*middle*), 5 µm (*right*). Images are mainly derived from frontal cortex sections of cases p1 and p2 (C2 C3: underlying white matter). Except C2 C3, images represent MIPs of z-stacks. White color indicates overlap of markers. **(b)** GLUA2 (G7) was used for western blots with human cerebral cortex lysate (Protein Medley, 50 µg), exposure time was 30 min for GLUA2 and 30 s for β-actin. Unit of the ladder is kilodalton. The same antibody is shown in a synaptic staining on frontal cortex sections (case p2). Scale bar, 2 µm. **(c)** Triple-staining with neuronal, axonal, and synaptic markers in a hippocampal section (case p2). For better visualization CAMKIIA is shown twice, in combination with SMI-312 (*left*) and with SYP (*right*). For the latter, an inset of the CA is provided in higher magnification (scale bar, 20 µm). CA = *cornu ammonis*, DG = dentate gyrus, alv = alveus, hil = hilus. Scale bars, 500 µm. **(d)** Tissue block with striatum (I, II) and pallidum (III). Immunolabeled sections from this block show matrix (TH high) and striosome (TH low) compartments in the caudate nucleus (I). VGLUT1 expression is strong in putamen (II) and absent in pallidum (III). G5 images represent MIPs of z-stacks. Scale bars, 400 µm (TH), 10 µm (VGLUT1). **(e)** VGLUT1 expression in the superior frontal gyrus and white matter (*top*). Intensity profile of the dashed area (raw data), normalized (norm.) to the maximum intensity (*bottom*). The green line marks the gray/white matter border, which was approximated by the abrupt reduction of the VGLUT1 signal. a.u. = arbitrary unit. Scale bar, 150 µm. Sections in (d) and (e) were derived from cases p1 and p2. The upper-case letter at the top of each image refers to the entities outlined in Figure 4a and the combination of letter and number allows for identification of the applied antibody listed in Table 2. See also source data files.

**Figure 4-figure supplement 3.**
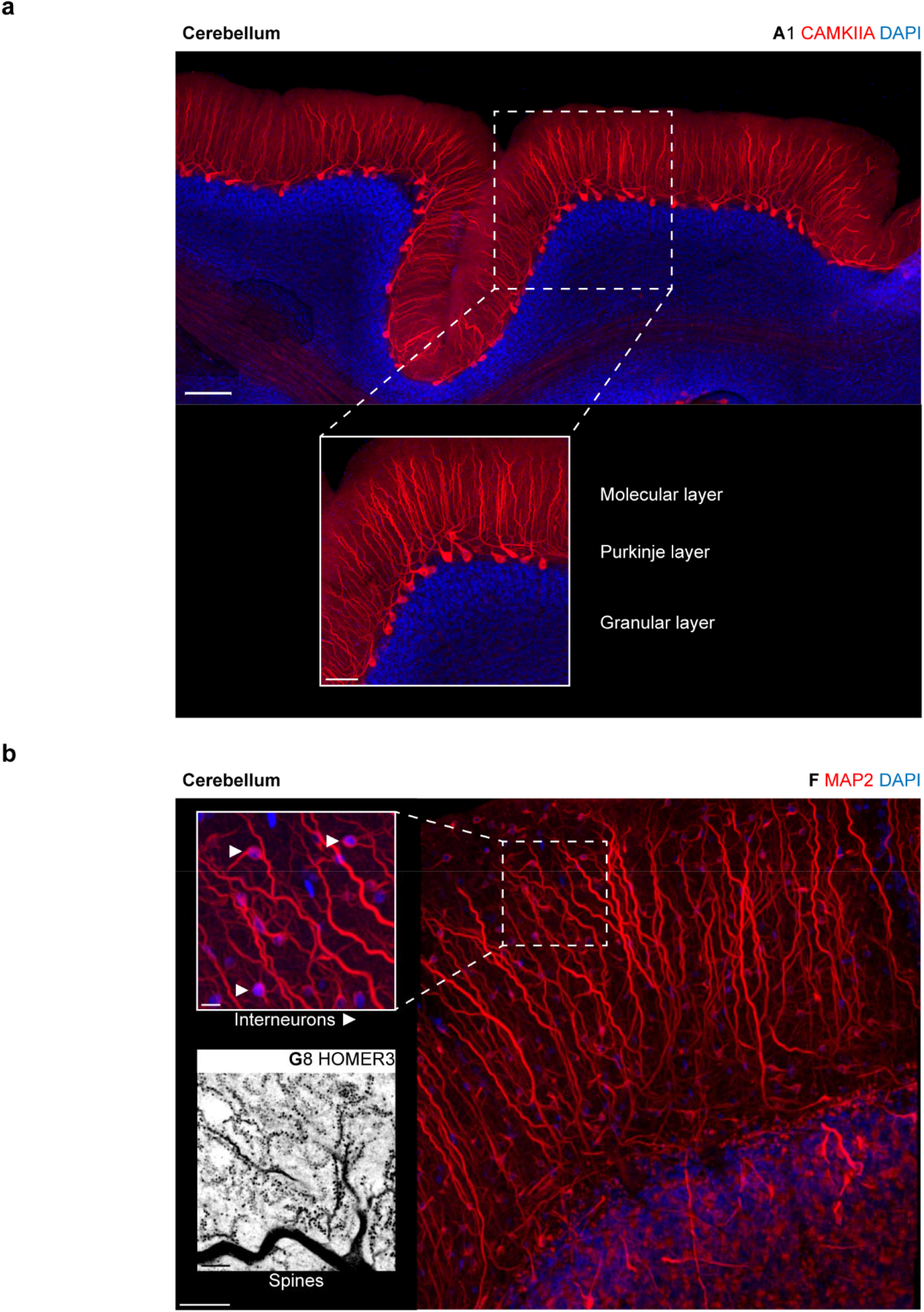
3D tilescans of neurocellular and synaptic markers in the human cerebellum after CLARITY. **(a)** CAMKIIA in the cerebellum, MIP of stack with x ∼3.2 mm, y ∼1.5 mm, and z = 54 µm. Scale bars, tilescan 200 µm, inset 100 µm. Stainings were performed on case p1. **(b)** 3D tilescan showing MAP2 and HOMER3 (shown as inverted image to highlight the spines of Purkinje cells in the molecular layer) in the human cerebellum, MIP of stack with z = 63 µm (MAP2). Scale bars, tilescan 50 µm, insets 10 µm (*top*), 5 µm (*bottom*).

**Figure 4-figure supplement 4.**
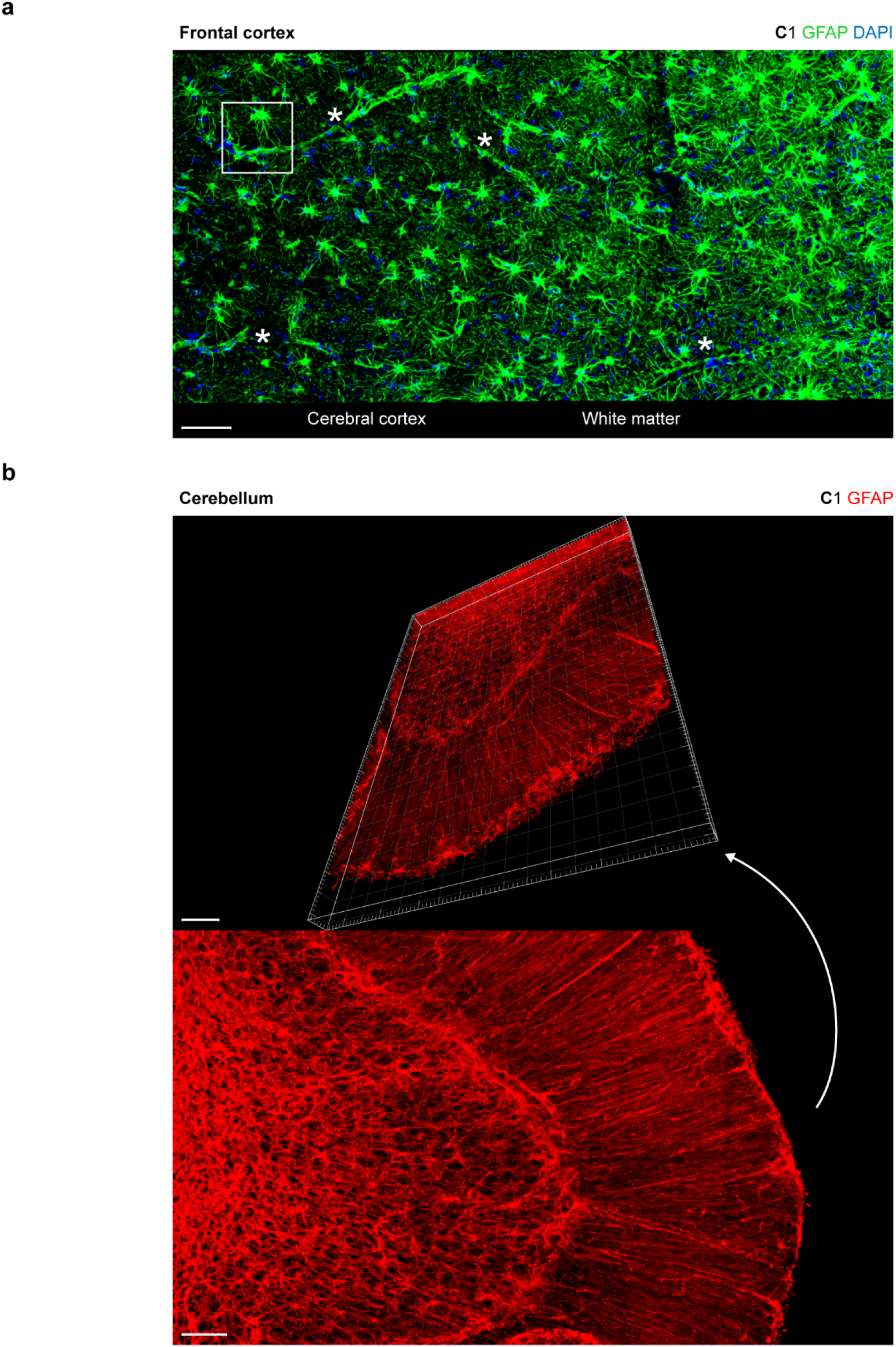
3D tilescans unveiling the extensive meshwork formed by GFAP-immunoreactive fibers after CLARITY. **(a)** Cerebral cortex and adjoining white matter stained for GFAP, MIP of stack with z = 10 µm is shown. Asterisks mark several sites where the blood-brain barrier can be recognized by GFAP-positive extensions wrapping the voids where a vessel runs along. The boxed area is shown in higher magnification in Figure 4b. Scale bar, 70 µm. **(b)** Cerebellar section stained with GFAP; dimensions in xyz, x = 848 µm, y = 1.1 mm, z = 56 µm. Scale bars, 100 µm (*top*), 70 µm (*bottom*). Sections were derived from perfusion-fixed cases (for (b) case p1).

**Figure 7-figure supplement 1.**
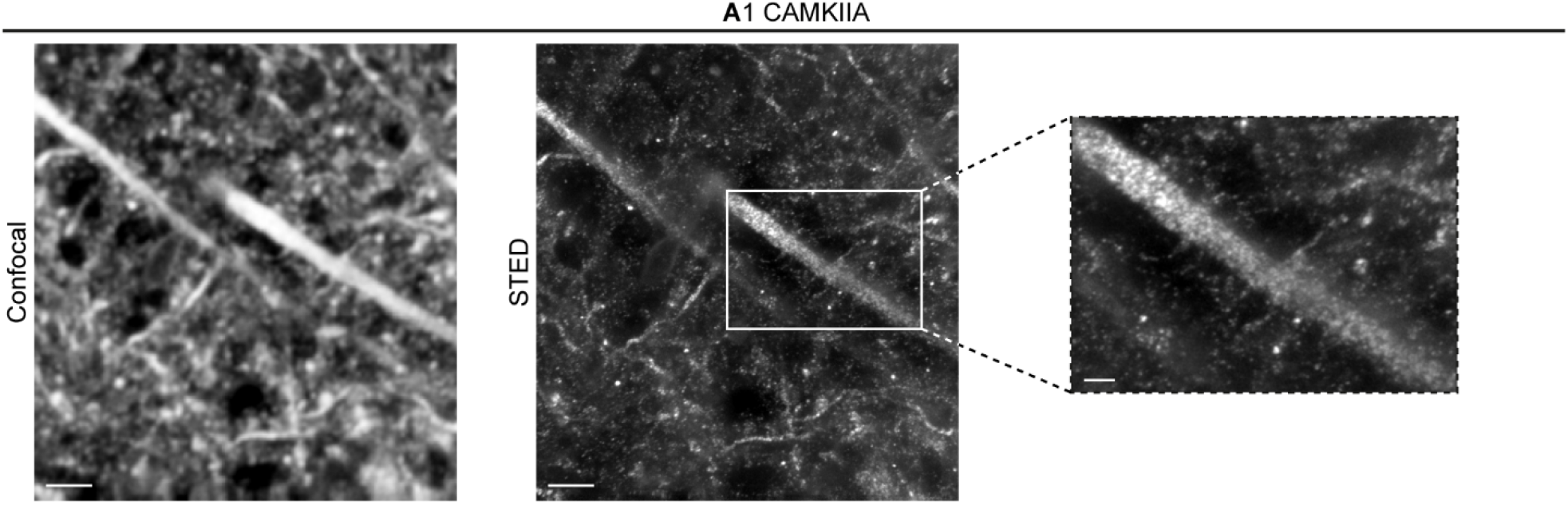
STED microscopy of CAMKIIA in human tissue after hCLARITY. CAMKIIA was acquired in confocal (*left*) and in STED mode (*middle and right*). Images are shown as MIPs of a stack with z = 3.6 µm. Z-drift was corrected with the Huygens Object Stabilizer (SVI). A higher magnification of a dendrite is shown on the right. Scale bars, 3 µm (*left and middle*), 1 µm (*right*). Pixel size for this confocal image is 100 nm, sections were incubated in hydrogel for seven days.

## 14. Rich Media Files: Videos

### Captions

**Video 1**

**3D imaging of AD tau pathology.** AT8-positive material (neurofibrillary tangles and neuropil threads) was stained in a 70 µm thick paraffin section derived from the cerebral cortex of an immersion-fixed case (LSM 710 microscope, Zeiss, 40x objective, NA 1.3, stack size 35.18 µm, step size 1.066 µm). Here, hydrogel embedding took place for 11 days, clearing time was 17 days. The section was mounted in FocusClear.

**Video 2**

**3D imaging of Aβ pathology (cored plaque) and VGLUT1.** Co-staining of a cerebral cortex section from a perfusion-fixed case with 4G8 (red) and a presynaptic marker (VGLUT1, G5, white) (confocal microscope, 40x objective, NA 1.15, stack size 24.77 µm, step size 0.36 µm).

**Video 3**

**3D imaging of Aβ pathology in vessels (CAA).** Immunostaining of a cerebral cortex section from an immersion-fixed case with 4G8 (confocal microscope, 63x objective, NA 1.3, stack size 39.64 µm, step size 0.28 µm).

**Video 4**

**3D imaging of GFAP-positive astrocytes and contribution to the blood brain barrier.** Immunostaining of white matter from a perfusion-fixed case with GFAP (C1, red), counterstaining with DAPI is shown in blue (confocal microscope, 40x objective, NA 1.15, stack size 98 µm, step size 1 µm).

**Video 5**

**3D imaging of MBP.** Immunostaining of a cerebral cortex section from case p2 with MBP (D3). The fibers were traced as surfaces within Imaris software and color-coded according to their z-position (confocal microscope, 40x objective, NA 1.15, stack size 63.53 µm, step size 0.5 µm).

**Video 6**

**3D imaging of MAP2-positive neurons and dendrites.** Immunostaining of a cerebral cortex section from case i4 with MAP2 (F). The video traces MAP2-positive cells and fibers from the outer cortex surface down to the beginning of the white matter (confocal microscope, 40x objective, NA 1.15, stack size 15.01 µm, step size 0.5 µm).

**Video 7**

**3D imaging of HOMER3 in the cerebellum.** HOMER3 (G8) is shown in red, DAPI counterstaining in blue, case p1 (confocal microscope, 40x objective, NA 1.15, stack size 47.52 µm, step size 0.22 µm).

**Video 8**

**3D imaging of collagen IV in the cerebral cortex.** Collagen IV (J1) is shown in red, DAPI counterstaining in blue, case p2 (confocal microscope, 40x objective, NA 1.15, stack size 46 µm, step size 1 µm).

**Video 9**

**3D imaging of smooth muscle actin (SMA) in the cerebral cortex.** SMA (J2) is shown in white, DAPI counterstaining in blue, perfusion-fixed case (confocal microscope, 63x objective, NA 1.3, stack size 34.27 µm, step size 0.35 µm).

**Video 10**

**3D super-resolution microscopy of CAMKIIA and VGLUT1 in the cerebral cortex.** CAMKIIA (magenta, A1, 1:200) and VGLUT1-ms (1:200, Synaptic Systems, Cat. No. 135511, RRID: AB_887879) were co-stained. Zoom-in at the end of the video highlights synapses (STED microscope, stack size 2 µm, step size 250 nm). Here, hydrogel embedding took place for 1 days, clearing time was 2 days. Z-planes were adjusted individually for brightness and contrast.

## 15. Reference to uploaded Source Data Files

Figure 2-Source Data (.xlsx)

Figure 2-figure supplement 1-Source Data (.xlsx)

Figure 4-figure supplement 2-Source Data (.xlsx)

Figure 4-figure supplement 2-Source Data (.zip folder)

Figure 7-Source Data (.xlsx)

